# A scRNA-seq atlas of chronic inflammatory skin diseases

**DOI:** 10.64898/2026.05.15.725328

**Authors:** Sabina Gansberger, Inigo Oyarzun, Martin Simon, Shawn Ziegler-Santos, Hao Yuan, Wolfgang Bauer, Philipp Tschandl, Wolfgang Weninger, Johanna Strobl, Sophie Frech, Maksim V. Plikus, Maria Kasper, Johannes Griss

## Abstract

Inflammatory skin diseases (ISDs) affect up to 25% of the global population. Yet, large-scale comparative single-cell RNA-sequencing (scRNA-seq) analyses between ISDs are still missing. Here, we integrated scRNA-seq datasets spanning 27 skin diseases from 50 studies, comprising over 2 million cells from 441 samples. Using the healthy skin cell atlas as reference, we could build a robust ISD atlas that enabled us to differentiate universal inflammatory signatures and disease-specific ones. This highlighted, for example, a shared gene program between keratinocytes in atopic dermatitis and parapsoriasis, not present in cutaneous T-cell lymphoma, confirms the plasticity of Th17 cells throughout ISDs, defines specific macrophage signatures in acne, and reveals a yet undescribed role of mural cells in ISDs. This demonstrates the power of the ISD atlas as a resource to resolve disease-specific immune mechanisms. The complete atlas is available through an interactive online portal at https://isd-atlas.derma.meduniwien.ac.at.

## Introduction

Inflammatory skin diseases (ISDs) are a heterogeneous group of conditions characterized by chronic inflammation of the skin^1^, collectively affecting approximately 20-25% of the population^2^. They arise from multiple etiologies, including dysfunctional immune cells, environmental factors, and genetic predisposition. The most frequent ISDs include atopic dermatitis (AD), psoriasis, alopecia areata (AA), hidradenitis suppurativa (HS) and lichen planus (LP)^2^. ISDs often profoundly impact a patient’s quality of life^2,3^, making early and precise treatment critical for effective disease management. However, differential diagnoses pose significant challenges. A key example is distinguishing early-stage cutaneous T-cell lymphoma (CTCL), a type of non-Hodgkin lymphoma from AD, characterised by eczema and pruritus^4^. Clinically and histologically, these diseases may appear very similar and there is currently no definitive diagnostic test^5^. This highlights the need for precise markers that can be used in routine diagnostics.

Beyond diagnostic challenges, most ISDs currently lack targeted therapies. While psoriasis benefits from mechanistically well-studied treatments^6^, most ISDs are treated with nonspecific therapies such as topical corticosteroids or broad anti-inflammatory agents^7^. Drugs approved for psoriasis were recently also approved for HS, yet show less efficacy^8^. While ISDs were historically classified by clinical and histological features^9^, some authors suggest redefining this taxonomy based on molecular profiling. Eyerich *et al.*^10^ categorized ISDs into immune modules based on dominant T cell responses: Th1 (AA, vitiligo), Th2 (AD, bullous pemphigoid (BP)), Th17 (psoriasis, HS), and Treg-associated (scleroderma, keloid). This framework more closely matches therapeutic strategies by highlighting shared immune pathways.

However, Th2-driven diseases like AD and BP illustrate the limitations of this immune-module classification: nemolizumab, an AD therapy, can paradoxically trigger BP^4^, revealing another layer of complexity. Moreover, emerging studies highlight that these diseases are not as homogenous as expected. Examples include acute AD (Th2-dominated) *vs.* chronic AD (Th1-shifted)^11^, psoriasis (Th17/Th1 variability)^12^, and HS (T cell-dependent/independent pathways)^13^. These complexities demand deeper molecular profiling to refine individual therapeutic strategies.

While bulk RNA-sequencing has uncovered disease-defining molecular abnormalities^7,14,15^, its inability to resolve cell type-specific signatures^16^ has hindered progress. Single-cell RNA-sequencing (scRNA-seq) offers unparalleled resolution but has largely been confined to intra-disease comparisons (lesional *vs.* nonlesional/healthy) and only few studies have comparatively explored multiple skin diseases^16,17^. To address this gap, we present an ISD atlas encompassing scRNA-seq data from 27 inflammatory skin conditions. The integration of multiple independent datasets per disease ensures the robustness of our resource and findings. This atlas reveals both shared and disease-specific molecular programs, while also exposing sample-preparation artifacts that could confound interpretation. Our findings underscore the potential of integrating large-scale scRNA-seq data to achieve a more comprehensive understanding of diverse pathologies of a single organ.

The atlas is publicly available under: https://isd-atlas.derma.meduniwien.ac.at as part of the Human Cell Atlas^18^.

## Results

### Integrating Multi-Source Data to Reveal Shared and Disease-Specific Cellular Signatures

To better understand the differences between ISDs we compiled a large collection of publicly available scRNA-seq datasets^17,19–59^, integrating them with inhouse data of previously published studies^60–63^ and novel data from BP and AA. This ISD atlas spans 27 diseases and healthy skin, originating from 50 publications, including 441 samples (diseased: 401; healthy: 40), with a total of 2,004,183 cells (Fig. 1a, Supplementary Fig. 1a). Metadata was extracted from the original publications and harmonized (Supplementary Table 1, Supplementary Fig. 1b). To achieve detailed and robust cell annotations, we utilised marker genes from the healthy skin cell atlas (HSCA) (Extended Data1). Following first level clustering we identified T cells, natural killer cells, B cells, plasma cells, macrophages, dendritic cells, plasmacytoid dendritic cells (pDC), mast cells, endothelial and lymphatic endothelial cells, melanocytes, Schwann cells, fibroblasts, mural cells, keratinocytes and sweat gland cells (Fig. 1b). T cells, fibroblasts and keratinocytes were the largest cell populations in the dataset (536k, 374k and 359k cells, respectively) (Fig. 1b, Supplementary Fig. 1c). This comprehensive atlas enables systematic exploration of shared and disease-specific features.

**Figure 1:**
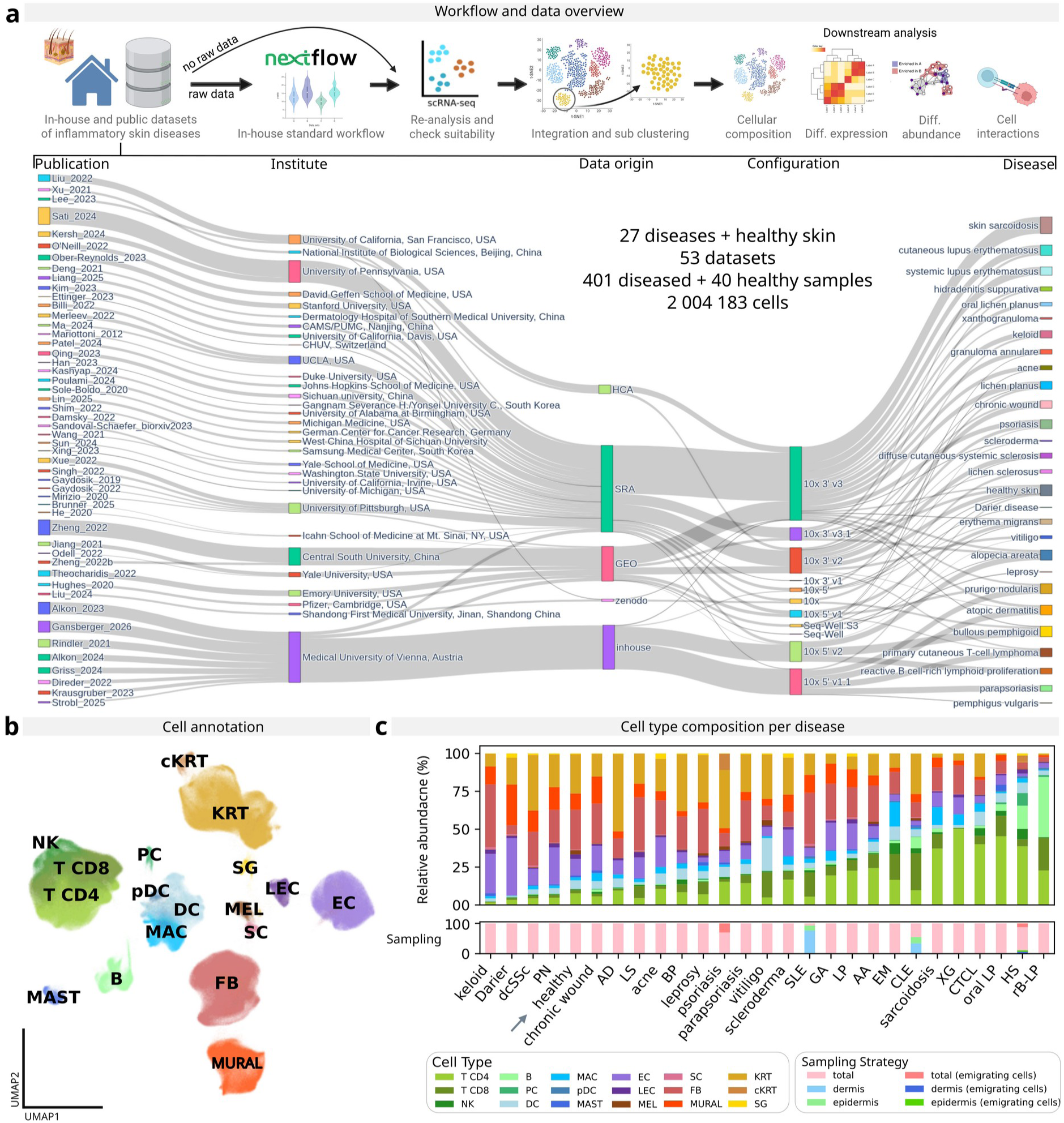
Dataset overview and overall cellular composition of ISDs. **a)** Study Workflow: Overview of the analytical workflow (Created in BioRender. Griss, J. (2026)) and details of the included studies. **b)** Integrated Cell Type Annotation: Uniform manifold approximation and projection (UMAP) embedding of the integrated dataset, annotated by cell type. **c)** Cellular Composition by Disease: Relative abundance of cell types (top) and sampling strategy (bottom) across diseases. Note: Samples containing CD45+-sorted cells were excluded from analyses in panels c) B, B cells; DC, dendritic cells; EC, endothelial cells; FB, fibroblasts; KRT, keratinocytes; cKRT, corneum Keratinocytes; LEC, lymphatic endothelial cells; MAC, macrophages; MAST, mast cells; MEL, melanocytes; MURAL, mural cells; NK, natural killer cells; pDC, plasmacytoid dendritic cells; PC plasma cells; SG, sweat glands: SC, schwann cells; AA, alopecia areata; AD, atopic dermatitis; BP, bullous pemphigoid; CLE, cutaneous lupus erythematosus; CTCL, primary cutaneous T-cell lymphoma; dcSSc, diffuse cutaneous systemic sclerosis; EM, erythema migrans; GA, granuloma anulare; HS, hidradenitis suppurativa; LP lichen planus; LS, lichen sclerosus; PN, prurigo nodularis; PV, pemphigus vulgaris; rB-LP, reactive B cell-rich lymphoid proliferation; SLE, systemic lupus erythematosus; XG, xanthogranuloma

While integrating multi-source datasets introduced technical challenges, such as biases stemming from sample preparation and processing (Extended Data 2), our analysis achieved robust harmonization (Supplementary Fig. 2). Dataset-specific variation primarily reflected intrinsic biological heterogeneity, with minimal methodological artefacts.

Analysis of cellular composition across ISDs revealed disease specific immune signatures. While most ISDs exhibited T cell expansion, this was absent in keloids, Darier disease, diffuse cutaneous systemic sclerosis (dcSSc), and prurigo nodularis (PN). B and plasma cells uniquely dominated the infiltrate in reactive B cell rich lymphoid proliferation (rB-LP), a non-malignant B lymphoid proliferating disorder^64^, and in HS, characterised by painful abscesses in intertriginous regions^65^, matching previous reports^61,66^. Macrophages were most prevalent in cutaneous sarcoidosis, a granulomatous disorder^48^, and dendritic cells in vitiligo, in-line with known vitiligo pathomechanisms^48,67^. Keratinocyte proportions exceeded 50% in AD, psoriasis, and BP, independent of body site (Extended Data 2), matching previous reports^68,69^. Relative fibroblast abundance was reduced compared to healthy skin in all diseases except keloid, likely due to immune cell infiltration in all ISDs but the latter (Fig. 1c). These shifts underscore key cellular drivers of ISD pathogenesis and confirm our resource’s ability to recapitulate known disease pathomechanisms.

### Disease specific overview and biases

To distinguish universal from disease-specific inflammatory responses, we quantified differentially expressed genes (DEGs) in each cell type relative to healthy controls (Fig. 2a, Supplementary Table 2). Fibroblasts, keratinocytes, and endothelial cells consistently showed high universal DEG counts across ISDs (detailed below), highlighting that comparisons between healthy and diseased samples in a single ISD often identify nonspecific, universally upregulated genes. Analysis of uniquely upregulated genes revealed several disease specific cellular drivers (Fig. 2a, Supplementary Table 3): NK cells in CTCL, dendritic cells and melanocytes in vitiligo and Schwann cells in keloids, aligning with known pathogenic drivers^40,67,70^. Moreover, mural cells showed specific signatures in systemic lupus erythematosus (SLE). Although their potential role in inflammation is increasingly recognized^71^, their precise functional contribution in SLE remains unclear. This demonstrates the atlas’ ability to distinguish between universal and disease-specific cellular changes.

**Figure 2:**
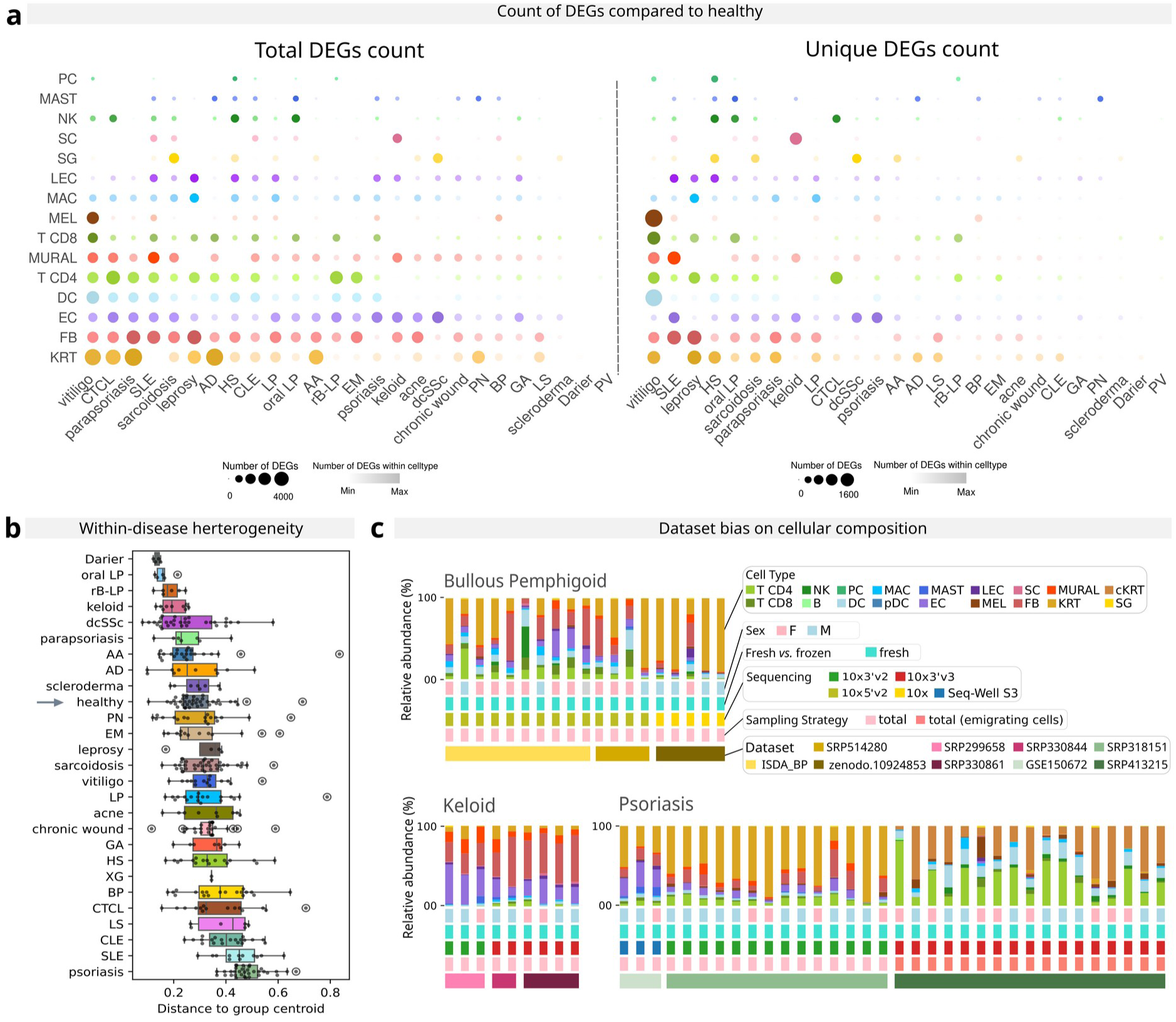
Disease specific overview and biases. **a)** Differential Gene Expression Analysis: Left: Number of total differentially expressed genes (DEGs) in each disease compared to healthy samples. Right: Number of unique DEGs in each disease compared to healthy samples. Dotsize represents the number of DEGs, color is scaled within cell type in order to highlight in which disease the cell type showed the highest number of DEGs and y axis are ordered by total number of DEGs per disease. **b)** Cellular Composition Diversity: Diversity index of cellular composition differences between samples within each disease. Outliers are indicated by circles. **c)** Dataset Bias: Stacked bar plots showing the relative frequency per sample, grouped by disease and stratified as follows (top to bottom): Cell type composition, Sex distribution, Fresh vs. frozen sample status, Sequencing technology, Sampling strategy. Note: Samples containing CD45+-sorted cells were excluded from analyses in panels in b) and c). B, B cells; DC, dendritic cells; EC, endothelial cells; FB, fibroblasts; KRT, keratinocytes; cKRT, corneum Keratinocytes; LEC, lymphatic endothelial cells; MAC, macrophages; MAST, mast cells; MEL, melanocytes; MURAL, mural cells; NK, natural killer cells; pDC, plasmacytoid dendritic cells; PC plasma cells; SG, sweat glands: SC, schwann cells; AA, alopecia areata; AD, atopic dermatitis; BP, bullous pemphigoid; CLE, cutaneous lupus erythematosus; CTCL, primary cutaneous T-cell lymphoma; dcSSc, diffuse cutaneous systemic sclerosis; EM, erythema migrans; GA, granuloma anulare; HS, hidradenitis suppurativa; LP lichen planus; LS, lichen sclerosus; PN, prurigo nodularis; PV, pemphigus vulgaris; rB-LP, reactive B cell-rich lymphoid proliferation; SLE, systemic lupus erythematosus; XG, xanthogranuloma

Certain ISDs exhibit large inter-patient heterogeneity^22,72^ complicating therapeutic strategies. To quantify this variability, we computed within-disease dispersion based on relative cell type frequencies using the Bray-Curtis distance (Fig. 2b). This metric quantifies the dissimilarity in species composition and assigns values to each sample ranging from 0 (identical to group mean) to 1 (completely dissimilar). Most ISDs showed substantial heterogeneity, such as CTCL or BP (Fig. 2b). Heterogeneity in psoriasis is likely due to differences in sample preparations (Fig. 2c, Extended Data 2). However, keloid and dcSSc remained relatively homogenous across multiple independent datasets (Fig. 2b,c). These findings underscore a disease specific biological variability independent of technical batch effects.

### T and NK cells show shared and unique features across ISDs

T cells are central to inflammation and tissue repair^73^, making them key therapeutic targets^74^. To find disease specific differences, we subclustered 575,046 T and NK cells to arrive at a detailed classification, including various helper (Th; *CD4*+), cytotoxic (Tct; *CD8*+) and innate lymphoid cells (ILCs; *CD3*-) subtypes, (Fig 3a, Supplementary Fig. 3a) and calculated the relative composition per disease (Fig 3b). AD, BP and PV had the highest Th2 frequency. PV simultaneously had the highest Th17 signature, which matches prior observation that Th17 cells play an important role in this antibody-mediated disease^75^. Chronic wounds also showed high Th17 signatures (matching previous reports^76^), as well as the classical Th17 diseases psoriasis and HS (Supplementary Fig. 3b)^10^. NK cells were enriched in HS and chronic wounds. Overall, ISDs could be grouped depending on their dominant T cell subset. For example, LS and oral LP were very high in CD8 Tct1 cells, AD and psoriasis in T central memory (Tcm), and sarcoidosis and leprosy in Th1 (Fig 3b). This implies the existence of shared mechanisms across certain ISDs that have not yet been investigated. All together, these findings highlight that our atlas is able to recapitulate established knowledge on ISDs while offering new insights on shared T cells pathomechanisms.

**Figure 3:**
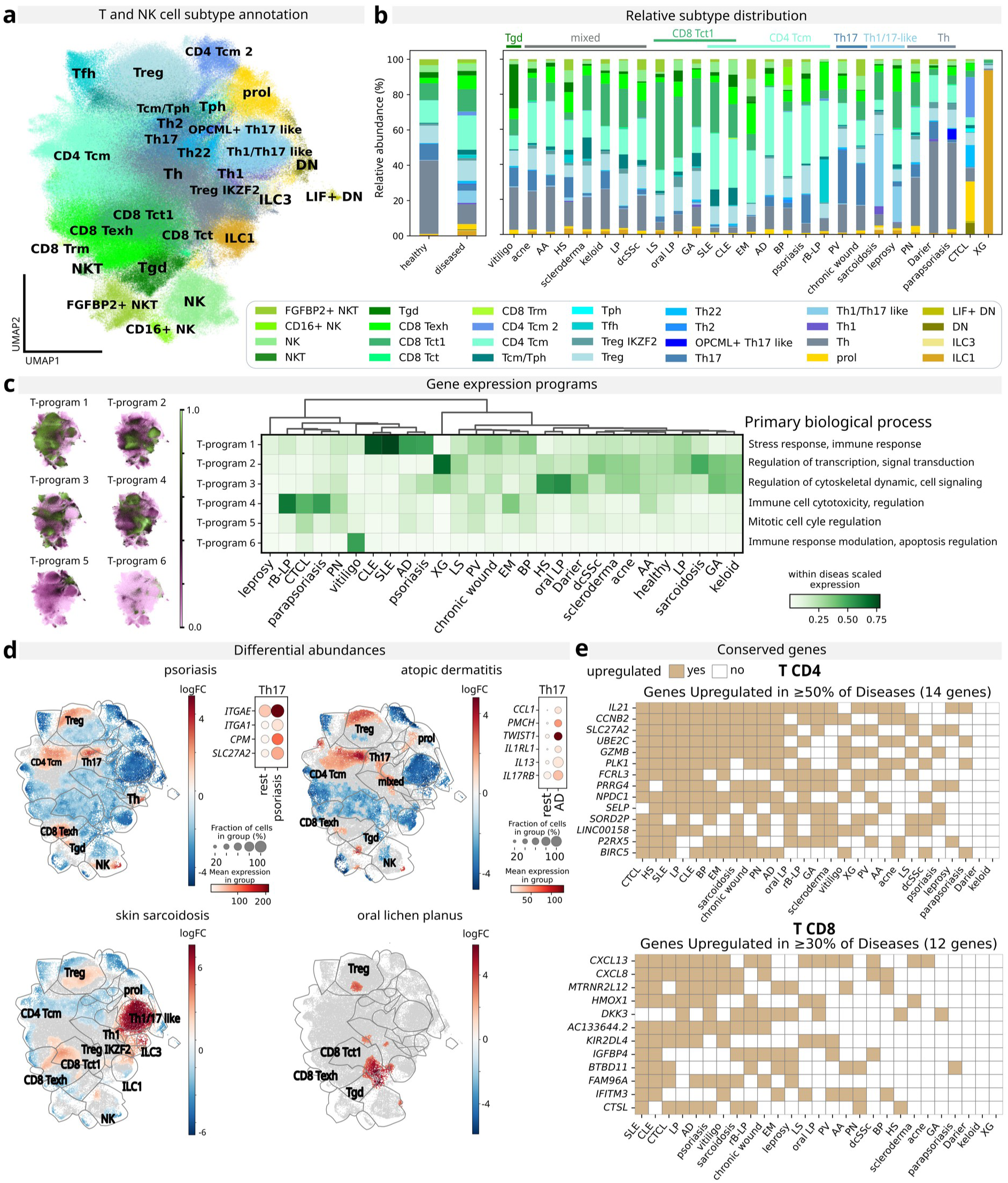
T and NK cells show shared and unique features across ISDs. **a)** T and NK Cell Subtype Annotation: UMAP embedding illustrating the annotation of T and NK cell subtypes. **b)** T Cell Subset Distribution: Relative abundance of T cell subsets in healthy samples and diseased samples (mean), and stratified by disease. ISDs are grouped by similar cell type distribution patterns. **c)** Gene Expression Programs: UMAP plot and heatmap of gene expression T-programs identified using cMNF. **d)** Differential Abundance of Neighborhoods and Pseudobulk DEG Expression: Differential abundance testing for selected diseases. Neighborhoods are colored by log fold change, non-differential abundance neighborhoods (FDR 10%) are colored grey. Dotplot of pseudobulk expression of DEGs (log₂ fold change > 1, adjusted p-value < 0.05) in highlighted populations. **e)** Conserved Upregulated Genes in Disease: Top percentage of conserved upregulated genes in disease compared to healthy controls (log₂ fold change > 2, adjusted p-value < 0.5). DN, double negative T cells; gdT, gamma-delta T cells; ILC, innate lymphoid cell; NK, natural killer cells; NKT, natural killer T cells; prol, proliferating cells; Tcm, central memory T cells; Texh, exhausted T cells; Tct, cytotoxic T cells; Tfh, follicular T helper cells; Th, T helper cells, Tph, peripheral T helper cells; Treg, regulatory T helper cells; Trm, tissue resident memory T cells

To find shared gene expression programs across ISDs, we applied consensus non-negative matrix factorization (cNMF)^77^, identifying six distinct modules (Fig 3c, Supplementary Table 4). T-program 1, characterised by interferon and MHC class I genes, was enriched in CLE, SLE, AD and psoriasis. The co-expression of genes associated with T-program 1 in T cells could be validated in public Xenium data from AD^78^ (Supplementary Fig. 3c). The top associated pathways included “Interferon Signaling” and “Antigen processing” (Supplementary Fig. 3d). Differential abundance testing within diseases revealed similar profiles for AD and psoriasis, including a higher Th17 abundance (Fig 3d, Supplementary Fig. 3e). Psoriatic Th17 cells upregulated tissue resident marker genes such as *ITGAE* and *ITGA1*, compared to Th17 cells of all other ISDs. While AD is a Th2 driven disease, Th17 cells in AD lesions are associated with disease severity^79^. AD’s Th17 cells showed upregulation of *CCL1*, *TWIST1* but also of *IL1RL1* and *IL13* (typically associated with Th2 cells) (Supplementary Table 5). This could indicate that a subset of Th2 cells interacts with Th17 cells, based on their expression of *IL17RB*, or a presence of plastic Th17/Th2 cells^80,81^, which have been reported to induce IgE production in asthma^81^. Nevertheless, differentiating these possibilities necessitates further investigation, yet it underscores the atlas’ potential to uncover novel pathomechanisms.

T-program 2 (enriched in sarcoidosis and xanthogranuloma) featured phosphodiesterase genes (Fig. 3c, Supplementary Table 4), which are emerging targets in granuloma treatment^82,83^. Sarcoidosis presented an increased differential abundance in the Th1/17-like population (Fig. 3d, Supplementary Fig. 3e) (*IFNG*+, *TNF*+, *RORC*+, *CCR6*+ but *IL17−*, as previously described^84^), supporting its classification as a Th17.1 disorder^23,85^. This cluster was universally conserved across sarcoidosis datasets (Supplementary Fig. 4a). Salt-inducible kinase (SIK)-genes, linked to anti-fibrotic effects^86^, were also upregulated in T-program 2, matching common grouping of fibrotic diseases (dcSSc, scleroderma and keloid). These findings underscore T-program 2’s role in granulomatous and fibrotic ISDs.

HS and oral LP had the highest expression in T-program 3, which was defined among others by *DOCK* and *PKC,* which are key regulators of immune response^87,88^, but have not been directly associated with these diseases yet. Oral LP showed higher abundance of CD8 T cells with increased histone genes (Supplementary Table 5). The highest expression in T-program 4, associated with immune cell activation and cytotoxic response, was found in rB-LP, CTCL and parapsoriasis, which share clinical features and are associated with lymphocyte proliferation. T-program 5 (cell cycle regulation; Supplementary Fig. 3d) had the highest expression in CTCL, matching known T cell expansion^89^. Cell-cell interaction mapping uncovered CTCL-specific immune crosstalk: *CD4*+ T–NK cell interactions via *MHCI–KIR* (NK inhibition^90^) and *PPIA–BSG* (pro-inflammatory/tumorigenic^91^), with the latter lacking direct CTCL evidence and *KIR3DL2* being an emerging drug target of CTCL^92^. Unlike other ISDs, CTCL exhibited no C-type lectin–*KLRB1* signaling highlighting disease-specific immune dynamics (Supplementary Fig. 4b). Overall, this data highlights the potential for identifying yet unrecognised similarities between ISDs.

To identify conserved upregulated genes, we analyzed DEGs in T cells across ISDs compared to healthy skin (p_adj < 0.05, LFC > 2). The top conserved upregulated genes were mainly associated with immune response and cell adhesion functions (Supplementary Table 6). In CD4+ T cells universal upregulation of *IL21* (differentiation and proliferation of lymphocytes^93^), *FCRL3* (Treg suppression^94^) and *PLK1, CCNB2, UBE2C* (cell cycle regulators^95–97^) was observed (Fig. 3e). CD8 T cells exhibited less conservation, with only *CXCL13* (also known as B-lymphocyte chemoattractant^98^), upregulated in more than 50% of ISDs. Contrary to that, clear B cell enrichment was only observed in HS and rB-LP (Fig. 1c), suggesting that *CXCL13* in most ISDs recruits T cells and/or dendritic cells via *CXCR5*^99^ rather than B cells. Also *CXCL8*, which recruits and activates neutrophils^100^, *HMOX1* and *DKK3*, which can influence the immune response^101,102^, were among the top upregulated genes. Shared natural killer cell genes included *ADGRG3* (cell-cell interactions^103^), *KIR2DL4* (inhibitory receptor^90^) and *ITGA1* (cell migration^104^) (Supplementary Fig. 5c, Supplementary Table 4). This highlights the atlas’ strength at identifying universally upregulated genes, enabling a focus on disease-specific features in downstream analyses.

### Fibroblasts express with the highest number of conserved genes upon inflammation

Fibroblasts play a pivotal role in maintaining skin architecture and rapidly produce and remodel extracellular matrix (ECM) following tissue injury, while also modulating immune responses through cytokine secretion^105,106^. Subclustering 374,763 fibroblasts identified 13 different clusters, including fibroblastic reticular cell-like (FRC-like; *CH25H*+) subtypes, hair follicle associated (HF-as; *TNMD*+) populations, Schwann cell like (*SCN7A*+), reticular (universal fibroblasts; *PI16*+) and papillary (superficial fibroblasts; *COL13A1*+) subtypes (Fig. 4a, Supplementary Fig. 5a). Notably, two myofibroblast (MyoFB; *ADAM12*+) clusters were exclusively detected in diseased samples, matching prior reports^78^ (Fig. 4b). Relative cell type distribution per disease revealed three dominant patterns: (1) MyoFB-associated: characterized by disease associated MyoFBs (keloid, chronic wound, HS); (2) FRC-like-dominated (SLE, CLE, EM); and (3) healthy-like (psoriasis, AD) (Fig. 4b). These findings highlight the functional plasticity of fibroblasts in inflammatory skin diseases.

**Figure 4:**
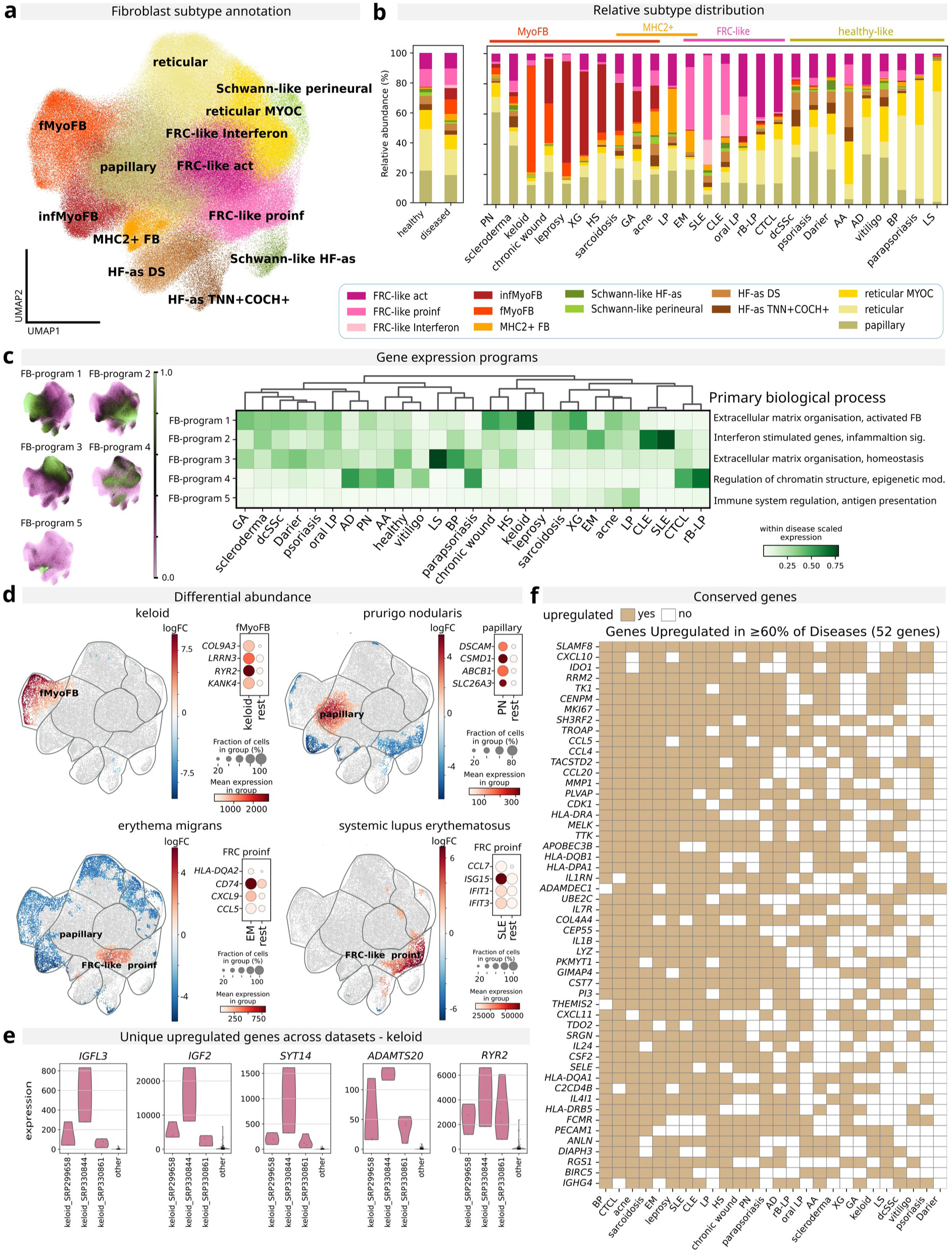
Fibroblasts express with the highest number of conserved genes upon inflammation. **a)** Fibroblast Subtype Annotation: UMAP embedding illustrating the annotation of fibroblast subtypes. **b)** Fibroblast Subset Distribution: Relative abundance of fibroblast subsets in healthy samples and diseased samples (mean), and stratified by disease. ISDs are grouped by similar cell type distribution patterns. **c)** Gene Expression Programs in Fibroblasts: UMAP plot and heatmap of gene expression FB-programs identified using cMNF. **d)** Differential Abundance of Neighborhoods and Pseudobulk DEG Expression: Differential abundance testing for selected diseases. Neighborhoods are colored by log fold change, non-differential abundance neighborhoods (FDR 10%) are colored grey. Dotplot of pseudobulk expression of DEGs (log₂ fold change > 1, adjusted p-value < 0.05) in highlighted populations. **e)** Conserved Upregulated Genes in Fibroblasts: Pseudobulk expression of uniquely upregulated genes conserved across all keloid datasets compared to all other diseases (log₂ fold change > 2, adjusted p-value < 0.05). **f)** Conserved Upregulated Genes in Disease vs. Healthy: Top percentage of conserved upregulated genes in disease compared to healthy controls (log₂ fold change > 2, adjusted p-value < 0.05). fMyoFB, fascia-like myofibroblasts; FRC-like, fibroblastic reticular cell-like; HF-as DS, hair follicle associated dermal sheet; infMyoFB, inflammatory myofibroblasts;

Our data showed that HS, chronic wounds and keloid share a distinct gene program (FB-program 1; Fig. 4c), characterised by collagens and other ECM associated genes, with association towards “ECM organisation” and collagen pathways (Supplementary Table 4, Supplementary Fig. 5b). HS specifically exhibited inflammatory MyoFB enrichment. Previous studies showed that lesional HS fibroblasts specifically upregulate *CXCL13* and *SFRP4* compared to healthy skin^107^. While *SFRP4* is upregulated in various ISD fibroblasts, high *CXCL13* expression is highly specific for HS (Supplementary Fig. 6c). This demonstrates the atlas’ utility in refining disease-specific molecular signatures.

Chronic wounds and keloids exhibit distinct fibroblast profiles and are dominated by fascia-like MyoFB (fMyoFB) (Fig. 4b), marked by *ACTA2, COL3A1, COL5A1*, and enhanced ECM/TGF-β signaling (vs. inflammatory MyoFB^78^). Keloids, characterized by persistent fibroblast activation leading to pathological scarring^108^, showed a significantly higher abundance of fMyoFB (Fig. 4d, Supplementary Fig. 5d) with unique upregulation of ECM remodeling (*LOXL3, LRRN3, PTGS1)*, and neural genes (*RYR2, GABRA5, VSNL1, OPCML*), as well as *KANK4* (Supplementary Table 5), matching prior reports^109^. Additionally, *SYT14, ADAMTS20, RYR2*, genes not yet associated with keloid (Fig. 4e, Supplementary Table 7), were consistently upregulated between datasets, similar to IGFL3 and IGF2 which are already linked to keloid pathogenesis^110^, highlighting the unique features of keloid fibroblasts.

FB-program 3 (ECM organization) differed from FB-program 1 by expressing homeostatic dermal fibroblast markers (*CD34, PI16, DCN*) rather than activated ECM-producing fibroblasts (*FAP, POSTN, COL11A1*) and was enriched in LS and BP. Key pathways included “neutrophil degranulation”, and complement genes (*CFD/H*), aligning with BP’s blister pathogenesis^111,112^. FB-program 5 (antigen presentation) showed the highest expression in LP, fitting the observed increase of *MHCII* FB, which has not been described previously (Fig. 4b,c, Supplementary Fig. 6d). This underlines the functional adaptivity of fibroblasts across ISDs.

FRC-like fibroblasts are similar to FRC found in lymphoid organs and support recruitment of immune cells and inflammatory signalling within the skin^106^. SLE, CLE and erythema migrans (EM) showed the highest relative frequency of proinflammatory FRC-like fibroblasts and the highest expression in gene FB-program 2, which is associated with interferon and inflammation signalling (Fig. 4c, Supplementary Fig. 5b). Differential abundance testing highlighted distinct subsets of proinflammatory FRC-like fibroblasts in EM and SLE (Fig. 4d, Supplementary Fig. 5d). In EM, these cells upregulated *CXCL9* and *CCL5*, which are associated with T cell recruitment^54,113^ as well as *MHCII* and *CD74*, which are involved in antigen presentation^114^. In contrast, SLE FRC-like proinflammatory cells upregulated genes associated with interferon signalling, matching previous reports of high interferon signatures^106^ (Supplementary Table 5). This highlights that our approach is able to detect even subtle differences within specific cellular subsets.

PN, characterized by the formation of intensely itchy nodules and dysregulated fibroblasts^49^, showed enriched abundance of papillary fibroblasts. Compared to other diseases, this cell population showed an increase in *DSCAM* and *CSMD1*, which are involved in neuronal development^115,116^ and *ABCB1* and *SLC26A3* which are involved in signalling regulation^117,118^ (Fig. 4c, Supplementary Table 5). None of these genes have been associated with PN in the past, but could explain the neural dysregulation in PN^119^.

Across ISDs, fibroblasts exhibited the greatest number of DEGs compared to healthy skin (Fig. 1e), underscoring their universal transcriptional response to inflammation. Analysis of conserved gene expression revealed 128 genes consistently upregulated in over 50% of ISDs (Supplementary Table 6), with enriched pathways including “signaling by interleukins” and “MHC class II antigen presentation” (Supplementary Fig. 5e). Notably, 55 genes, including chemokines (*CXCL10, CCL4, CCL5, CCL20*), cell cycle regulators (*MKI67, CDK1, RRM2*^120^), and ECM remodeling enzymes (*MMP1*), were upregulated in over 60% of ISDs. *SLAMF8*, the most frequently upregulated gene, is critical for fibroblast migration, proliferation, and inflammation^121^ (Fig. 4f). These findings emphasize the conserved and central role of fibroblasts in ISD pathogenesis.

### Schwann cells feature prominently in keloid

Schwann cells are known to be major contributors to keloid formation^40^. Subclustering of Schwann cells led to three distinct clusters: myelinating Schwann cells (*MBP*+, *MPZ*+), non-myeloating Schwann cells (*NCAM1*+, *SCN7A*+) and a cluster nearly exclusively originating from keloid samples and featuring upregulated *IGFBP5*, *PTN*, as previously described^40^, as well as *KIRREL3,* which is connected to synapse formation^122^ (Supplementary Figure 6a-c). Difference between Schwann cells from keloid and other ISDs is reflected in their high number of unique DEGs (Fig. 1d, Supplementary Table 3). In total, 753 genes were uniquely upregulated in keloid Schwann cells, including functions such as ECM remodeling, cell cycle progression and RHO GTPase effectors, which are key pathways in fibrosis^108^ (Supplementary Figure 6d).

### Mural cells emerge as new players in ISDs

Mural cells, consisting of pericytes and smooth muscle cells, are abundant in skin, but are typically not considered key drivers of ISDs. We observed that in HS, leprosy, chronic wounds and PN there is an increased abundance of *POSTN+* pericytes (Supplementary Figure 6e-g). In fibroblasts, increased expression of *POSTN* has been connected to PN previously^49^, but the changes in pericytes have not yet been investigated. Differential abundance testing confirmed the increase in *POSTN*+ pericytes in PN. CLE and SLE showed an expansion in *HAS2*+ pericytes (with role in ECM synthesis^123^) and *NR4A2*+ pericytes (known for regulating gene expression in response to inflammation^124^; Supplementary Figure 6g). Intriguingly, *NR4A2* drug targets are used in chronic autoimmune diseases like rheumatoid arthritis^125^, indicating that mural cells may play an active role in these ISDs.

### Most ISDs are dominated by the shared keratinocyte gene program

Keratinocytes, primary epidermal cells, maintain barrier integrity and modulate immunity via cytokine production^126^. Subclustering of 379,824 keratinocytes identified basal (*KRT15*+), spinous (*MT1G*+), granular (*KRT10*+), and terminal (*ACER1*+) subsets, alongside disease-enriched populations such as hyperproliferative keratinocytes, which are marked by stress response genes (*KRT6A/B/C* and *KRT16*) and known to be associated with wound healing and inflammation^127^. This cell population was dominant in chronic wounds (as previously described^128^), LS, and PN (Fig. 5a,b, Supplementary Figure 8a). Based on the cellular composition, ISDs can be grouped into: (1) differentiated KRT dominated; (2) hyperproliferating and *MHCII+* KRT associated; and (3) adnexal-associated diseases.

**Figure 5:**
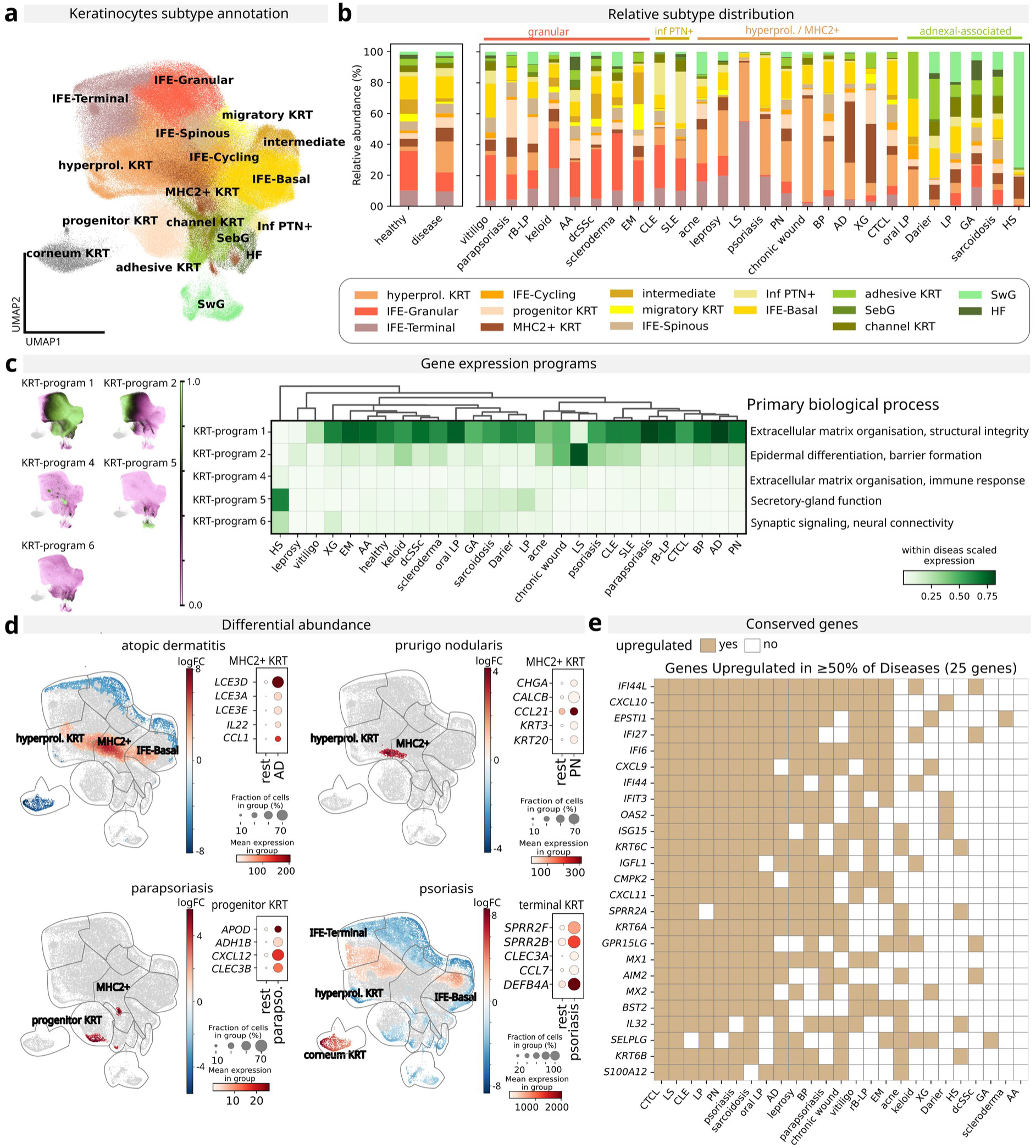
Most ISDs are dominated by the shared keratinocyte gene program. **a)** Keratinocyte Subtype Annotation: UMAP embedding illustrating the annotation of keratinocyte subtypes. **b)** Keratinocyte Subset Distribution: Relative abundance of keratinocyte subsets in healthy samples and diseased samples (mean), and stratified by disease. ISDs are grouped by similar cell type distribution patterns. **c)** Gene Expression Programs in Keratinocytes: UMAP plot and heatmap of gene expression KRT-programs identified using cMNF. **d)** Differential Abundance of Neighborhoods and Pseudobulk DEG Expression: Differential abundance testing for selected diseases. Neighborhoods are colored by log fold change, non-differential abundance neighborhoods (FDR 10%) are colored grey. Dotplot of pseudobulk expression of DEGs (log₂ fold change > 1, adjusted p-value < 0.05) in highlighted populations. **e)** Conserved Upregulated Genes in Disease: Top percentage of conserved upregulated genes in disease compared to healthy controls (log₂ fold change > 2, adjusted p-value < 0.05). cKRT, corneum keratinocytes; HF, hair follicle, SebG, sebaceous gland cells; SwG, sweat gland cells

Six keratinocyte gene programs were identified, with KRT-program 1 enriched in the majority of ISDs (Fig. 5c). The top ranking pathways associated with this KRT-program were “ECM organisation”, “laminin interactions” and “collagen formation” (Supplementary Figure 7c). In particular, AD and parapsoriasis showed prominent dominance of this KRT-program. AD presented with an increased abundance in *MHCII+* keratinocytes (Fig. 5d, Supplementary Figure 7d). These upregulated late cornified envelope genes (*LCE3D/A/E*) and immune modulators (*IL22*, *CCL1*; Supplementary Table 5). PN similarly showed *MHCII+*/hyperproliferative enrichment, but uniquely upregulated *KRT20/3* and *CHGA*, *CALCB* (neural signaling^129,130^), potentially linked to pruritus (Fig. 5d). Parapsoriasis featured progenitor keratinocytes expressing oxidative stress genes (*APOD*^131^, *ADH1B*^132^) as well as *CLEC3B* and *CXCL12*. Psoriasis showed an increased abundance in various keratinocyte populations, with terminally differentiated cells upregulating *SPRR2F/B, CLEC3A*, *CCL7*, and *DEFB4A* (immune modulation). KRT-program 2 (terminal differentiation) was enriched in LS, consistent with the known basal cell depletion^133^. HS, which was dominated by sweat gland cells (known hallmark of HS^134^), showed an increase in KRT-program 5 (secretory gland function). Overall, our analysis indicates that keratinocytes behave similarly in most ISDs with only minor disease specific changes.

We observed a conserved upregulation of 25 genes in more than 50% of ISD, including interferon-associated genes (*IFI44L*, *IFI27*, *IFI6*, *IFI44*), and chemokines (*CXCL9/10/11*). Moreover, stress response genes^127^ (*KRT6A/B/C*), which are key markers of hyperproliferative keratinocytes, were shared across ISDs, highlighting their important role in cutaneous inflammation and emphasizing keratinocytes’ dual structural and innate immune roles (Fig. 5e).

### Myeloid cells feature shared inflammatory mechanism between granulomatous diseases and acne

Myeloid cells, particularly macrophages, orchestrate inflammation through pro-and anti-inflammatory cytokine secretion and play a key role in multiple ISDs^135^. To dissect myeloid cell heterogeneity, we subclustered 189,789 macrophages, dendritic cells and pDC and arrived at a detailed classification including: neutrophils (NP; *FCGR3B*+), conventional dendritic cells (*CLEC9A*+ cDC1, *CLEC10A*+ cDC2), mature DC (matDC; *LAMP3*+), Langerhans cells (*CD207*+), monocyte-derived dendritic cells (moDC; *FCN1*/*ICAM*+), lipid-associated macrophages (LAMs; *TREM2*+, *APOE*+), perivascular macrophages (PVM; *SELENOP*+) and interstitial macrophages (IM; *CXCL3*+, *THBS1*+). Based on relative cell type abundance, ISDs could be grouped into: (1) NP-associated (HS, LS), (2) PVM-rich (scleroderma, EM), (3) pDC-rich (rB-LP), (4) LAM-rich (GA, sarcoidosis), (5) LC-associated (vitiligo), (6) matDC-rich (psoriasis) and (7) healthy-like (Figure 6a,b, Supplementary Fig. 8a).

**Figure 6:**
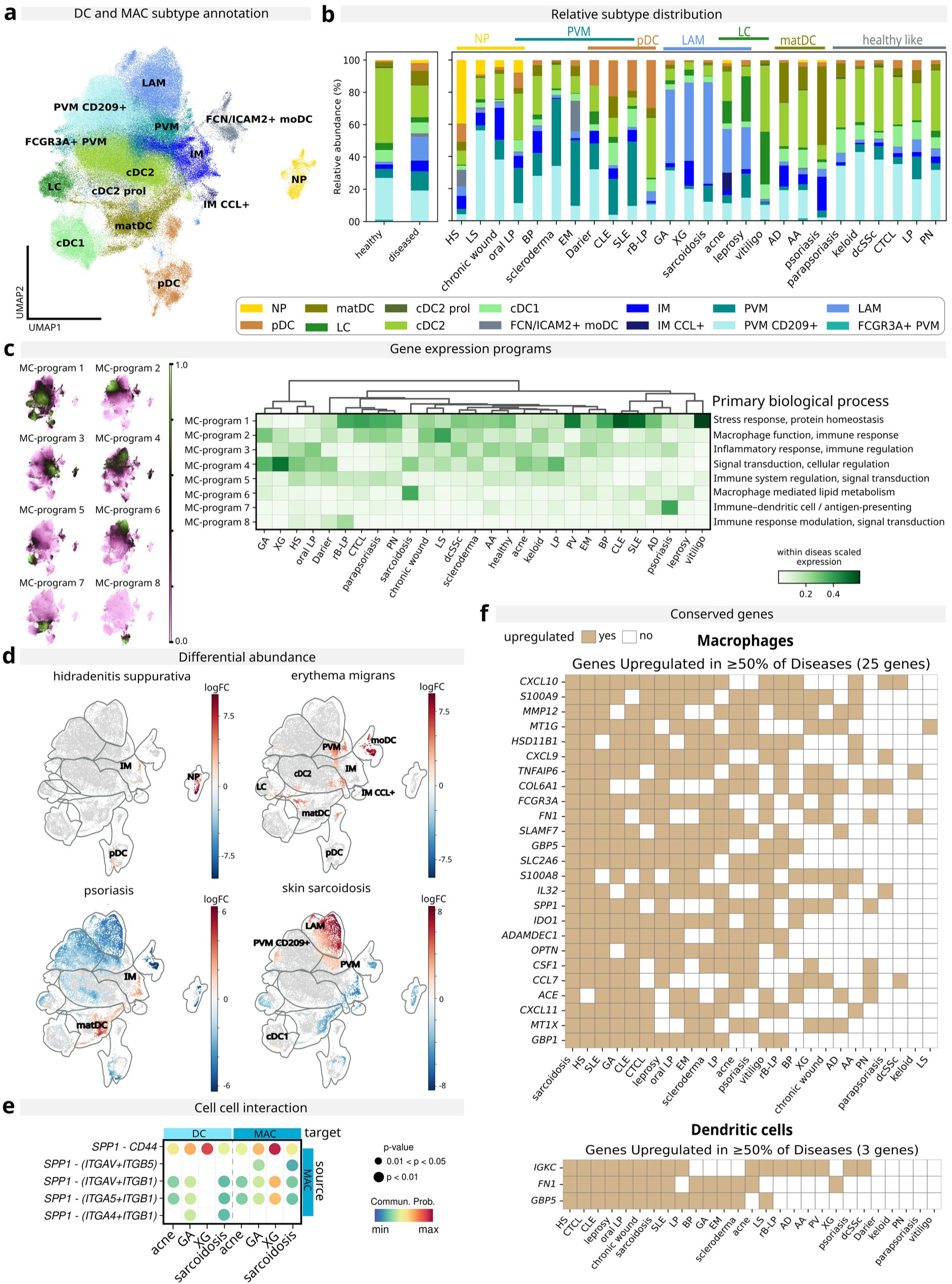
Myeloid cells feature shared inflammatory mechanism between granulomatous diseases and acne. **a)** Myeloid Cell Subtype Annotation: UMAP embedding illustrating the annotation of myeloid cell subtypes. **b)** Myeloid Subset Distribution: Relative abundance of myeloid subsets in healthy samples and diseased samples (mean), and stratified by disease. ISDs are grouped by similar cell type distribution patterns. **c)** Gene Expression Programs in Myeloid Cells: UMAP plot and heatmap of gene expression MC-programs identified using cMNF. **d)** Differential Abundance of Neighborhoods: Differential abundance testing for selected diseases. Neighborhoods are colored by log fold change, non-differential abundance neighborhoods (FDR 10%) are colored grey. **e)** Cell–Cell Interaction Network: Bubble plot of selected cell–cell interactions involving myeloid subsets. Note: negative values for all other ISDs were removed. **f)** Conserved Upregulated Genes in Disease: Top percentage of conserved upregulated genes in disease compared to healthy controls (log₂ fold change > 2, adjusted p-value < 0.05). cDC, conventional dendritic cells; IM, interstitial macrophage; LAM, lipid-associated macrophages; LC; matDC, mature dendritic cells; MC, myeloid cells; pDC, plasmacytoid dendritic cell; PVM, perivascular macrophage

LAMs were enriched in granulomatous diseases, specifically sarcoidosis (63%), XG (49%), GA (45%) and leprosy (26%) (Fig. 6b, Supplementary Fig. 8b). This aligns with prior reports of lipid metabolism in sarcoidosis^136^, dyslipidemia in GA^137^, and APOE upregulation in leprosy^138^. Granulomatous diseases also exhibited strong upregulation in MC-program 4 (signals transduction and cellular regulation; Fig. 6c). Notably, acne (27% LAMs) shared this signature, matching previous reports of *TREM2*+ macrophages in acne^139^ and suggesting mechanistic overlap between granulomatous ISDs and acne. Differential abundance analysis confirmed LAM enrichment in sarcoidosis, GA and acne (Fig. 6d, Supplementary Fig. 8c). Compared to other granulomatous diseases, sarcoidosis showed additional enrichment in MC-program 6 (macrophage mediated lipid metabolism), with “Neutrophil degranulation” as the top pathway (Fig. 6c, Supplementary Fig. 8d). Though NP are not classically linked to cutaneous sarcoidosis, their association with pulmonary sarcoidosis severity^140^ suggests potential systemic immune overlap. Cell-cell interaction mapping revealed unique macrophage-dendritic cell crosstalk in granulomatous diseases and acne, mediated by *SPP1-CD44*/integrin signaling, a pathway linked to lymphocyte recruitment^48^ but mainly described in cancer context^141^. While *SPP1* upregulation in acne’s myeloid compartment is documented^142^, its shared role in acne and granulomatous disease pathogenesis has not been recognized (Fig. 6e). These findings underscore the central role of macrophages in sarcoidosis and suggest a shared inflammatory mechanism with acne.

Besides granulomatous diseases, several additional ISDs showed unique signatures in the myeloid compartment. CLE, SLE and PV showed dominance in MC-program 1 (stress response), with “Interferon signaling” being the dominant pathway, matching previous reports for lupus^106^ (Supplementary Fig. 8d). MC-program 3 (interleukin signaling; Supplementary Fig. 8d) was enriched in various ISDs, indicating a universal immune response. In HS, a higher abundance of NP was observed (Fig. 6d, Supplementary Fig. 8c), consistent with their pathogenic role via NET release^143^. Erythema migrans exhibited moDC and PVM enrichment (Fig. 6d, Supplementary Fig. 8c), which has not been described previously, and a decrease in LC abundance, in concordance with recent findings^144^. Psoriasis showed matDC and IM expansion, alongside MC-program 7 (antigen presentation) upregulation (Fig. 6c,d, Supplementary Fig. 8c), featuring *CCL22* (Th2/Treg recruitment^145^), *IL23A/12B* (psoriatic relapse^146^), *IL15* (pathogenesis^147^), and *IL32* (cell-specific expression^148^) (Supplementary Table 4). This matches reports that *LAMP3*+ DCs are critical *IL23* producers that are enriched in psoriatic lesions^146^. Overall, our data highlights the disease specific features of myeloid cells in ISDs.

In addition to displaying disease specific features, myeloid cells also respond universally to inflammation. Comparative analysis revealed 25 genes universally upregulated in macrophages, irrespective of subtype, primarily linked to immune response and inflammation, including *S100A8/9*, *CXCL9/10/11*, *MMP12*, *CSF1*, *FCGR3A*, and *IL32*. By contrast, dendritic cells showed limited conserved MC-programs, with only three genes (*IGKC*, *FN1* and *GBP5*) consistently upregulated in >50% of ISDs (Fig. 6f, Supplementary Table 6). Notably, *IGKC* expression is usually not associated with dendritic cells and could hint towards a minor population of lymphoid origin dendritic cells (L-cDCs)^149^. Overall, these data highlight macrophage plasticity versus dendritic cell specificity in ISD pathogenesis.

### Endothelial cells overexpress Insulin receptor in chronic wounds

Endothelial cells, often overlooked in disease pathology, were prominent in Darier disease, chronic wounds, keloids, and PN (Fig. 1b). Subclustering of 205,324 endothelial cells identified an *INSR*-enriched cluster, significantly expanded in chronic wounds (Supplementary Fig. 9a,b). Endothelial cells are known contributors to wound healing^150^, but their direct role in wound healing efficacy is still unknown. *INSR* expression was the highest at wound edges and lowest in non-healers (healer vs nonhealer: Mann-Whitney U Test: p-value = 1.8e-19; Supplementary Fig. 9c). This was also found in a re-analysis of public Visium spatial transcriptomics data from acute wounds in different phases (Supplementary Fig. 9d)^151^. There, co-expression of PECAM1 and INSR was increased during the proliferative phase (Day 7 post-injury, Supplementary Fig. 9d). This aligns with murine studies of *Insr* expression during acute healing^152^ and a clinical data linking local insulin to wound repair^153^, demonstrating the potential of our atlas to reveal disease specific features that would stay undetected when comparing diseased to healthy skin only.

### Online resource of the inflammatory skin disease atlas

We created an online web resource using the vitessce framework^154^ that allows for easy and fast exploration of the inflammatory skin disease atlas (https://isd-atlas.derma.meduniwien.ac.at). Here, users can explore the main object as well as the different cell subclusters and plot expression levels of genes of interest per disease, dataset or customized groups (Fig. 7a,b).

**Figure 7:**
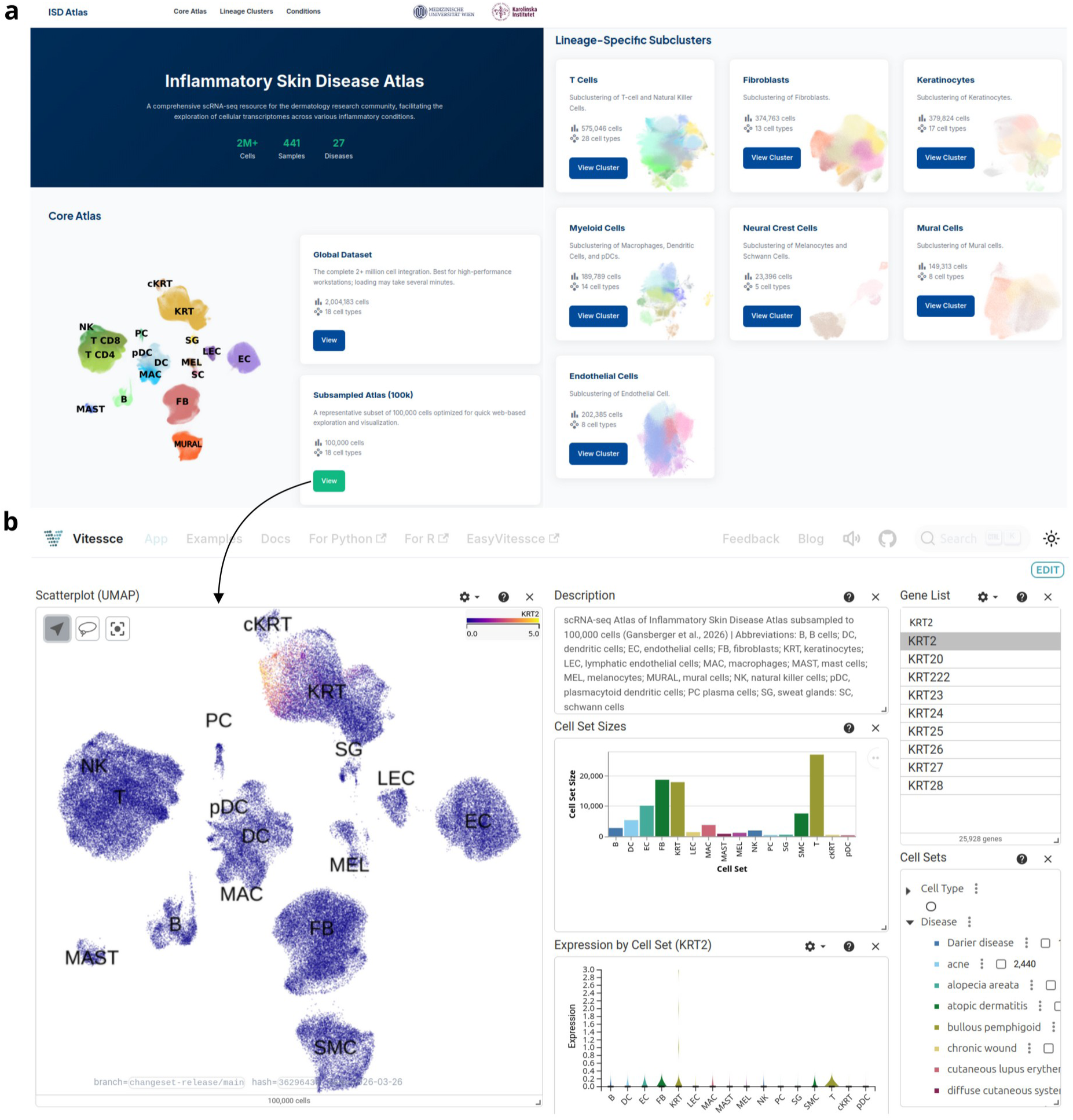
Online resource for easy and fast exploration of the Inflammatory Skin Disease atlas. **a)** Webtool Interface Overview: Users can select between exploring the main integrated dataset or specific subclusters for detailed analysis. **b)** Main Object Visualization: Example view of the main object, allowing users to color the UMAP embedding by cell type, disease, or dataset. The Cell Set Size panel displays the corresponding cell counts for each category. UMAP embedding showing KRT2 expression across all cells. The Expression by Cell Set panel highlights KRT2 expression per cell type.

## Discussion

Microarray^15^ and bulk RNA-seq studies^7,14^ of multiple ISDs have uncovered unique and shared inflammatory signatures. Nevertheless, scRNA-seq atlases of, for example cancers^155–157^, highlighted how this unprecedented resolution leads to novel insights into cellular heterogeneity and intercellular crosstalk. With respect to ISDs, studies comparing multiple diseases at single-cell level are still rare^16,17^. These datasets highlighted mutually exclusive features between distinct ISDs^138^ and specific properties of shared cell types across ISDs^158–160^, yet with certain limitations: (i) analysing only a few diseases^16,17^, (ii) using seqWell data^17^, which yields considerably fewer cells than 10X Genomics platforms^161^, or (iii) analysing only CD45+ cells, excluding full gamut of cells^16^. Thus, there remains an unmet need for a comprehensive single-cell atlas of diverse ISDs.

Here, we present the largest comparative scRNA-seq atlas of ISDs to date. Due to the large number of sequenced cells exceeding two million, their integration and comparison across diverse ISDs allowed us to achieve detailed annotation even among cell types with low abundance^162^, such as Schwann cells. Moreover, this atlas provides a comparative resource for both healthy tissue and clinically distinct inflammatory disease states. Using this atlas we were able to replicate many established findings both on cell abundance, for instance the presence of MyoFBs exclusively in diseased states^78^(Fig. 2b) and specific gene expression features, such as *CXCL13* overexpression in HS fibroblasts^107^ validating the robustness of our integrated dataset. Additionally, we were able to identify similarities between ISDs, such as upregulation of *CXCL13* in T cells (Fig. 2e) and *KRT6* in keratinocytes (Fig. 4e), as well as disease-specific features, including conserved fibroblast DEGs in keloid (Fig. 3e) and *POSTN*+ pericytes enrichment in prurigo nodularis (Supplementary Figure 6g), which have not been previously recognized. Moreover we found neutrophil degranulation enrichment in sarcoidosis (Supplementary Figure 8d), previously only linked to pulmonary sarcoidosis and a novel shared feature between acne and granuloma diseases (*SPP1* interactions in macrophages; Fig. 6e). Therefore, beyond confirming known biology, this atlas also reveals potentially novel disease-associated patterns.

While our atlas of ISDs provides valuable insights, several inherent challenges and limitations must be acknowledged. The availability of samples was constrained by pre-existing datasets, which were not always fully matched for patient sex, age and ethnicity, and exhibited unequal sample sizes across different conditions. Specifically, sample numbers were limited for conditions such as XG, PV, and Darier disease, which may affect the generalizability of our findings. Raw data were not available for all studies, which introduces additional noise, such as genome version bias (Extended Data 2). Additionally, the use of varying experimental protocols across independent studies introduced variability in cellular compositions. By harmonizing the datasets’ metadata, we clearly assessed the magnitude of these technical differences and demonstrated that our integration strategy still yielded robust results.

In conclusion, our data underscores the importance of integrating diverse datasets and diseases to refine our understanding of inflammatory skin diseases. Our atlas and online resource enable researchers to rapidly compare their findings across a broad range of datasets, serving as a foundation for future ISD research.

## Methods

### Data collection

Data from previously published inhouse studies as well as publicly available scRNA-seq data from inflammatory skin diseases were included (Supplementary Table 8). PubMed was searched for publicly available scRNA-seq datasets of dermatoses (excluding cancer, only exception was CTCL due to known challenges in differential diagnoses) and checked for suitability (E.g. number of samples). If the raw data was provided, SRA data was downloaded with the sra toolkit and converted to fastq files using fastq-dump (sra-tools^163^ version 3.1.0). If the raw files were provided as bam files, the files were converted to fastq using the bam2fastq^164^ tool. Otherwise the processed data was downloaded from GEO, zenodo or HCA respectively. For healthy control samples data from 7 different studies and different anatomical regions were included to reduce the batch effect as much as possible.

### Patient recruitment and sample processing of unpublished data

Skin punch biopsies from patients with alopecia areata and bullous pemphigoid were taken after written consent from patients recruited at the Medical University of Vienna, Austria (Ethics Committee vote: 1354/2021). Diagnosis was confirmed by a board-certified dermatopathologist in connection with routine clinical information.

Tissue samples were processed using the Dissociation Kit by Miltenyi Biotech (Bergisch Gladbach, Germany) and send for scRNA-seq using the Chromium Single Cell Controller with the Single Cell 5’ Library & Gel Bead Kit v2 (10X Genomics, Pleasanton, CA) following to the manufacturer’s protocol. Sequencing was performed in the 150bp paired-end configuration using the Illumina NovaSeq platform.

### Metadata harmonization

Metadata information was collected from the corresponding publication, and data deposition websites. The metadata was harmonized across the multiple studies through the usage of common variable names. This included among other disease description, sex, age, treatment and sample location of the individual samples as well as sequencing platform and original publication for the datasets.

### Data processing

The unpublished in-house data as well as all publicly available dataset where raw data was available were processed using 10x Genomics Cell Ranger v7.1.0^165^. The data was aligned to the human reference genome assembly “refdata-gex-GRCh38-2020-A”.

Standardized QC was performed on all samples (including datasets where only processed data was available), which included ambient RNA removal (DecontX^166^, celda_1.14.2), doublet removal (scDblFinder^167^, version 1.12.0), and filtering cells with a minimum of 500 genes, maximum of 20% mitochondrial genes, 5% haemoglobin and 1% platelet genes and genes expressed in less than 3 cells. For this R (version 4.1.0) and Seurat^168^ (version 5.0.1) were used. Additionally all samples were individually checked for quality and adjusted if necessary or excluded from the atlas.

Afterwards samples were converted to Scanpy AnnData objects and merged (scanpy^169^, version 1.11.4). Raw counts were normalized and log transformed. Highly variable genes, pca and k-nearest neighbor graphs were calculated before samples were integrated using the BBKNN algorithm (bbknn^170^, version 1.6.0), with sample_id as batch key.

Computation was performed on inhouse servers or on the Vienna Scientific cluster (vsc5) with a Scanpy docker image provided by gcfntnu, downloaded from dockerhub (docker://gcfntnu/scanpy; python version: 3.11.11, anndata==0.12.2, bbknn==1.6.0, scanpy==1.11.4). CellChat was run on vsc5 using the docker image provided by repbioinfo, downloaded from dockerhub (docker://repbioinfo/signacseuratcellchat:latest).

### Clustering and data annotation

To enable scalable clustering we aggregated the cells into so-called supercells using MiniBatch k-means (scikit-learn^171^, version 1.6.1) clustering algorithm. MinibatchKMeans was run with 1% of the cells with a batch size of 20,000. Average PCA representations for each supercell was calculated and used to construct a new AnnData object. Leiden clustering with a resolution of 2 and package’s implementation “igraph” was run on the supercell level AnnData object. Cluster assignments were mapped back to each corresponding cell in the full object.

Annotation of cell types was based on the human healthy skin atlas.

### Cell type composition within disease - relative frequency calculation and diversity index

To quantify compositional differences within the diseases Bray-Curtis dissimilarities^172^ were computed between the samples based on the relative cell type frequencies. For each disease centroids were calculated and the distance of each sample to the corresponding disease centroid was computed to obtain the within disease dispersion.

### Subcluster integrations

Cell types were extracted from the main object and the raw counts extracted. Each cell type object was integrated using the BBKNN algorithm. For integration of the T cells, Myeloid cells and endothelial cells ScVI^173^ (version 1.2.2.post2) algorithm was used with batch keys for raw_data, platform, dataset and sample_id to combat batch effects more effectively. The model was trained on 90% of data and 10% were used for validation. Cell type subtypes were annotated using canonical marker genes, the human healthy skin cell atlas reference and differentially expressed genes. Gene programs were identified using consensus NMF^174^ (cNMF, version 1.7.0) using 8.000 highly variable genes. Gene programs reflecting the genome bias (discussed in Extended Data 2) were excluded. Hu et al., Gene Set AI^175^ was used for discovery of gene set functions.

### Differentially expressed genes and differential abundance testing

DecoupleR package (version 2.0.7) was used to calculate pseudobulk per sample and cell type. EdgeR^176^ (pertpy^177^, version 0.10.0) was used for DEG analysis comparing each disease against healthy samples and filtering for FDR < 0,05 and LFC > 2. For dataset-conserved unique DEGs per disease and celltype, comparison was performed per dataset of target disease against all other diseases and only shared results were obtained.

Milo algorithm^178^ (pertpy^177^, version 0.10.0), was used to identify cell populations that showed a differential abundance in the individual diseases in the subclusters.

Enriched pathways were calculated using clusterProfiler^179^ (version 4.16.0) and REACTOME pathway database^180^ (version 1.52.0).

### Cell-cell communication

Cell-cell communication for each disease was computed using CellChat^181^ (version 2.2.0) and R (R version 4.4.3) and the human ligand-receptor interaction database (CellChatDB.human). Samples originating from dataset SRP413215 were excluded due to the observed influence of the sample preparation on the keratinocytes phenotype (Supplelementary Figure 2a). Cells expressing fewer than 200 genes as well as genes detected in less than 3 cells were excluded. CellChat was computed with the default parameters. Interactions with a minimum of 10 cells per interacting group were kept in order to remove low-confidence interactions. Interaction objects of all diseases were merged for comparative downstream analysis.

### Statistics and reproducibility

All data presented in this manuscript were derived from primary patient samples. All available samples were included in all analyses. No technical replicates were performed. All p-values reported in this manuscript were adjusted for multiple testing using the Benjamini–Hochberg false discovery rate (FDR) correction.

### Webtool

For easy and fast exploration of the atlas we used the framework provided by vitessce^154^ to create an online tool.

## Data Availability

Publicly available scRNA-seq dataset are listed in Supplementary Table 8. Newly generated sequencing data for AA and BP from this study are available from GEO (pending).

The integrated atlas and subclusters can be explored on an online web portal at: https://isd-atlas.derma.meduniwien.ac.at

The Python *AnnData* objects underlying the figures shown in this manuscript are available on zenodo under 10.5281/zenodo.20160794.

## Code Availability

No novel code was produced during this analysis.

## Abbreviations

AA: alopecia areata
AD: atopic dermatitis
BP: bullous pemphigoid
CLE: cutaneous lupus erythematosus
CTCL: primary cutaneous T-cell lymphoma
dcSSc: diffuse cutaneous systemic sclerosis
EM: erythema migrans
GA: granuloma anulare
HS: hidradenitis suppurativa
LP: lichen planus
LS: lichen sclerosus
PN: prurigo nodularis
PV: pemphigus vulgaris
rB-LP: reactive B cell-rich lymphoid proliferation
SLE: systemic lupus erythematosus
XG: xanthogranuloma

## Acknowledgements

This work was funded by research grants from the Austrian Science Fund (grant number P35937 and KLP7023723) and the LEO Foundation (grant number LF-OC-23-001394) to JG. IO was supported by a DOC fellowship from the Austrian Academy of Science (Grant number 27228). The Vienna Scientific Cluster (Project No. 71839) is gratefully acknowledged for providing computational resources. This publication is part of the Human Cell Atlas – www.humancellatlas.org/publications/

## Author Contributions

**Designed research** SG: JG **Sample acquisition** JG: WB: JS: SF **Sample analysis** MS: SZS: WW **Histopathological analysis** PT: WB **Data analysis** SG: IO: SF: HY: JG **Acquisition of funding** JG **Writing of manuscript** SG: MP: MK, JG All authors reviewed and approved the final version of the manuscript.

## Competing Interests

JG received personal fees from AbbVie: Eli Lilly: Pfizer: Boehringer Ingelheim and Novartis. JG is an investigator for Navigator Med. PT reports speaker fees from AbbVie and education project grants from Eli Lilly. JS received personal fees from Pfizer and Janssen.

All other authors declare no competing interests.

## Notes

https://isd-atlas.derma.meduniwien.ac.at

