## Supplementary information for "A scRNA-seq atlas of chronic inflammatory skin diseases"

**Key points:**

- diseased skin atlas, scRNA-seq, cutaneous inflammation, inflammatory skin diseases

**Abbreviations:** *AA, alopecia areata; AD, atopic dermatitis; BP, bullous pemphigoid; CLE, cutaneous lupus erythematosus; CTCL, primary cutaneous T-cell lymphoma; dcSSc, diffuse cutaneous systemic sclerosis; EM, erythema migrans; GA, granuloma anulare; HS, hidradenitis suppurativa; LP lichen planus; LS, lichen sclerosus; PN, prurigo nodularis; PV, pemphigus vulgaris; rB-LP, reactive B cell-rich lymphoid proliferation; SLE, systemic lupus erythematosus; XG, xanthogranuloma*

**Supplementary information** **1**

Supplementary Figure 1: Data overview 3

Supplementary Figure 2: Dataset density 4

Supplementary Figure 3: T and NK cells 6

Supplementary Figure 4: T and NK cells 8

Supplementary Figure 5: Fibroblasts 9

Supplementary Figure 6: Schwann cells and mural cells 10

Supplementary Figure 7: Keratinocytes 11

Supplementary Figure 8: Myeloid cells 12

Supplementary Figure 9: Endothelial cells 13

**Extended Data** **14**

Extended Data 1 - Harmonized cell annotation through the Healthy Skin Atlas 14

Extended Data Fig. 2: Mapping of inflammatory skin disease atlas to healthy skin cell atlas 15

Extended Data 2 - Technical bias through sampling and bioinformatics methods 16

Extended Data Fig. 2: Dataset bias based on data processing 18

Extended Data Fig. 3: Dataset bias based on sampling strategy 19

Extended Data only references 20

***Supplementary Figure 1: Data overview***

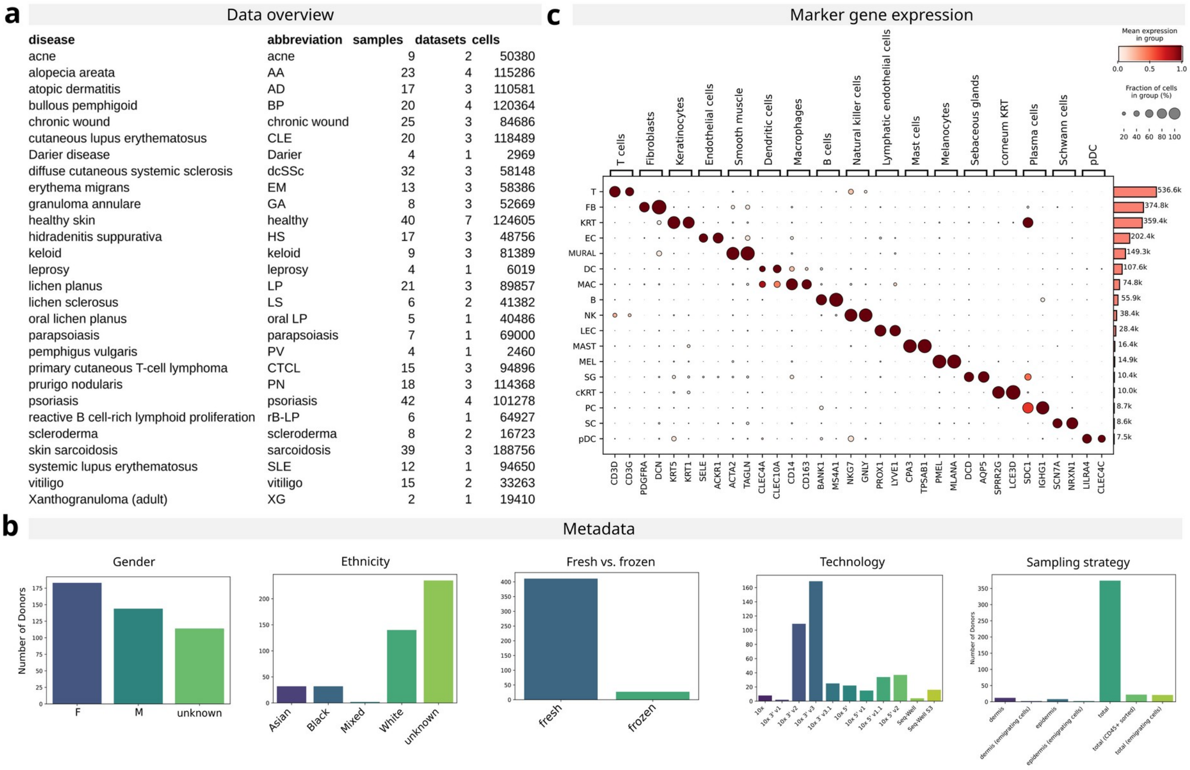

| disease | abbreviation | samples | datasets | cells |
|---|---|---|---|---|
| acne | acne | 9 | 2 | 50380 |
| alopecia areata | AA | 23 | 4 | 115286 |
| atopic dermatitis | AD | 17 | 3 | 110581 |
| bullous pemphigoid | BP | 20 | 4 | 120364 |
| chronic wound | chronic wound | 25 | 3 | 84686 |
| cutaneous lupus erythematosus | CLE | 20 | 3 | 118489 |
| Darier disease | Darier | 4 | 1 | 2969 |
| diffuse cutaneous systemic sclerosis | dcSSc | 32 | 3 | 58148 |
| erythema migrans | EM | 13 | 3 | 58386 |
| granuloma annulare | GA | 8 | 3 | 52669 |
| healthy skin | healthy | 40 | 7 | 124605 |
| hidradenitis suppurativa | HS | 17 | 3 | 48756 |
| keloid | keloid | 9 | 3 | 81389 |
| leprosy | leprosy | 4 | 1 | 6019 |
| lichen planus | LP | 21 | 3 | 89857 |
| lichen sclerosus | LS | 6 | 2 | 41382 |
| oral lichen planus | oral LP | 5 | 1 | 40486 |
| parapsoiasis | parapsoiasis | 7 | 1 | 69000 |
| pemphigus vulgaris | PV | 4 | 1 | 2460 |
| primary cutaneous T-cell lymphoma | CTCL | 15 | 3 | 94896 |
| prurigo nodularis | PN | 18 | 3 | 114368 |
| psoriasis | psoriasis | 42 | 4 | 101278 |
| reactive B cell-rich lymphoid proliferation | rB-LP | 6 | 1 | 64927 |
| scleroderma | scleroderma | 8 | 2 | 16723 |
| skin sarcoidosis | sarcoidosis | 39 | 3 | 188756 |
| systemic lupus erythematosus | SLE | 12 | 1 | 94650 |
| vitiligo | vitiligo | 15 | 2 | 33263 |
| Xanthogranuloma (adult) | XG | 2 | 1 | 19410 |



***a)*** *Data Summary: Overview of sample counts, datasets, and total cells stratified by disease.* ***b)*** *Metadata Summary: Comprehensive metadata profile of the included samples.* ***c)*** *Marker Gene Expression: Dot plot depicting marker gene expression, accompanied by bar plots indicating the corresponding cell type abundances.*

***Supplementary Figure 2: Dataset density***

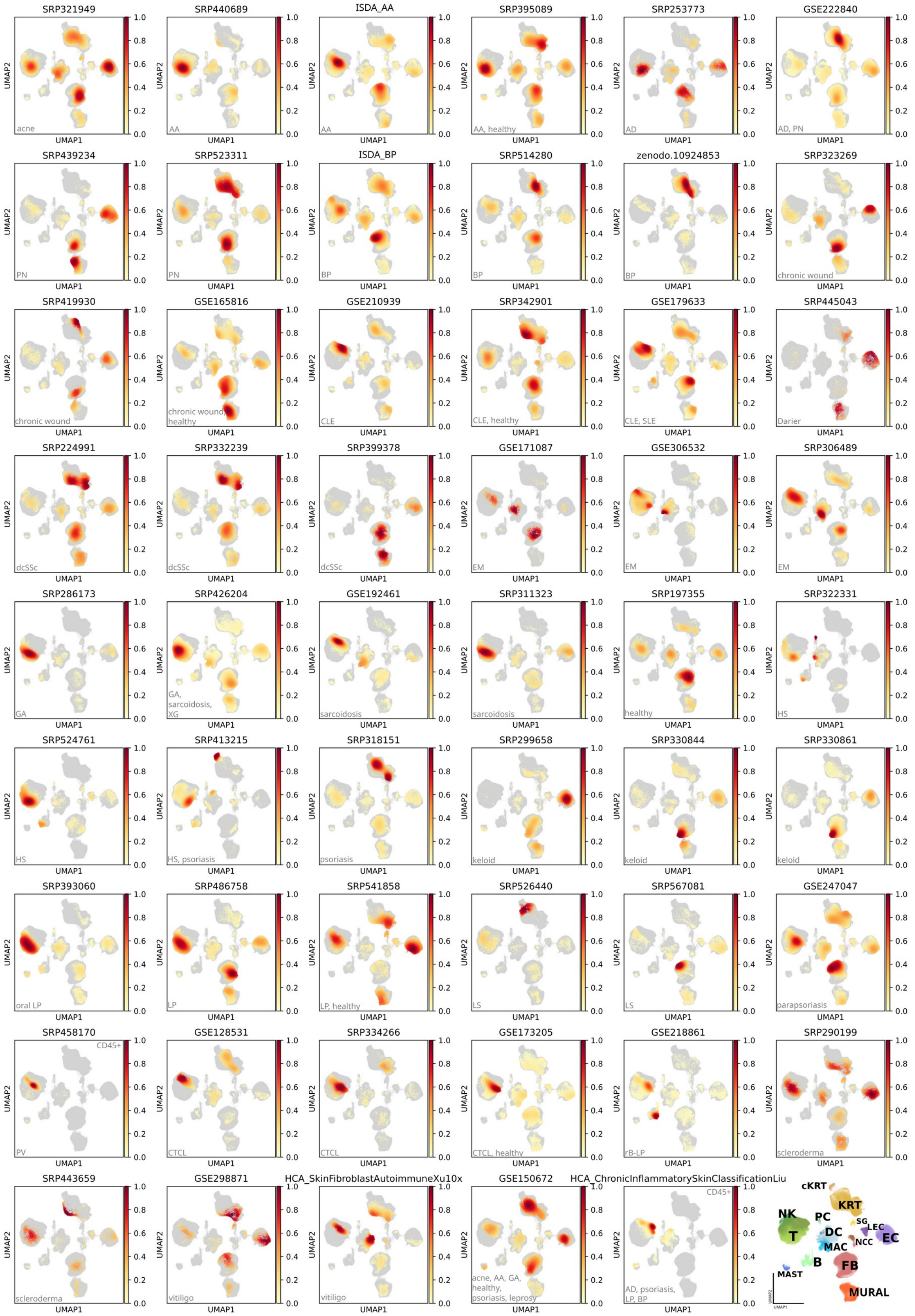


*Density distributions for each dataset are presented as individual plots. The associated disease for each dataset is indicated in the lower left corner of the respective plot. The final plot provides an overview of cell type annotations. B, B cells; DC, dendritic cells; EC, endothelial cells; FB, fibroblasts; KRT, keratinocytes; LEC, lymphatic endothelial cells;*

*MAC, macrophages; MAST, mast cells; MURAL, mural cells; NCC neural crest cells; NK, natural killer cells; PC plasma cells; SG, sweat glands*

***Supplementary Figure 3: T and NK cells***

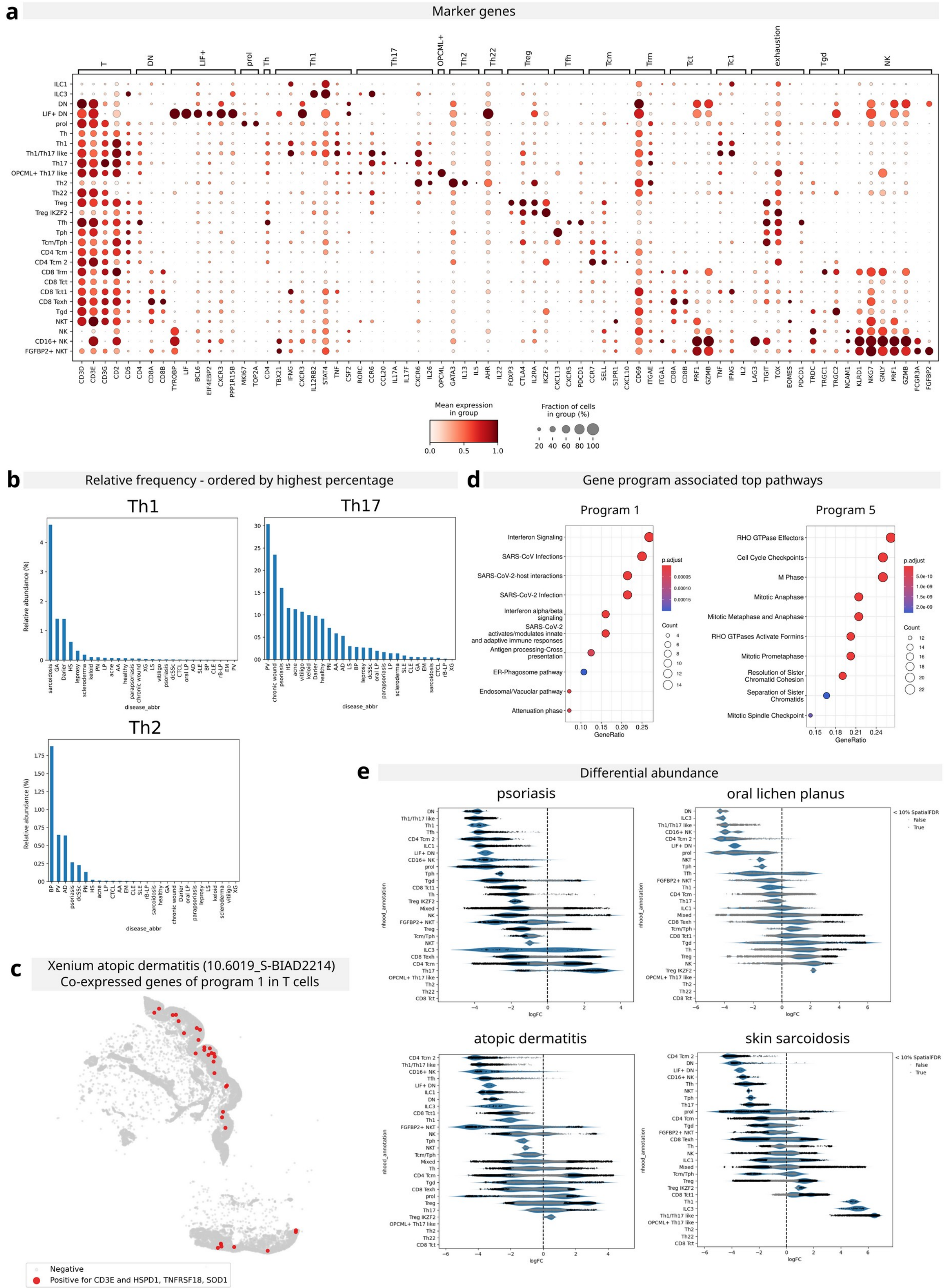


***a)** Marker Gene Expression: Dot plot illustrating the expression of marker genes across T and NK cell subtypes. **b)** Cell Type Frequency: Relative abundance of key T cell populations, stratified by disease. **c)** External Validation in Spatial Data (10.6019_S-BIAD2214): Xenium data of lesional atopic dermatitis to show co-expression of cNMF Program 1 genes in T cells: CD3E/HSPD1/TNFRSF19/SOD1 positive spots are highlighted in red. **d)** Pathway Enrichment: Top-ranked pathways associated with selected gene programs. **e)** Differential Abundance Analysis: Beeswarm plot of differential abundance* *testing for selected diseases. Each dot represents a neighborhood of cells, with significantly enriched neighborhoods (FDR < 10%) highlighted in black.*

***Supplementary Figure 4: T and NK cells***

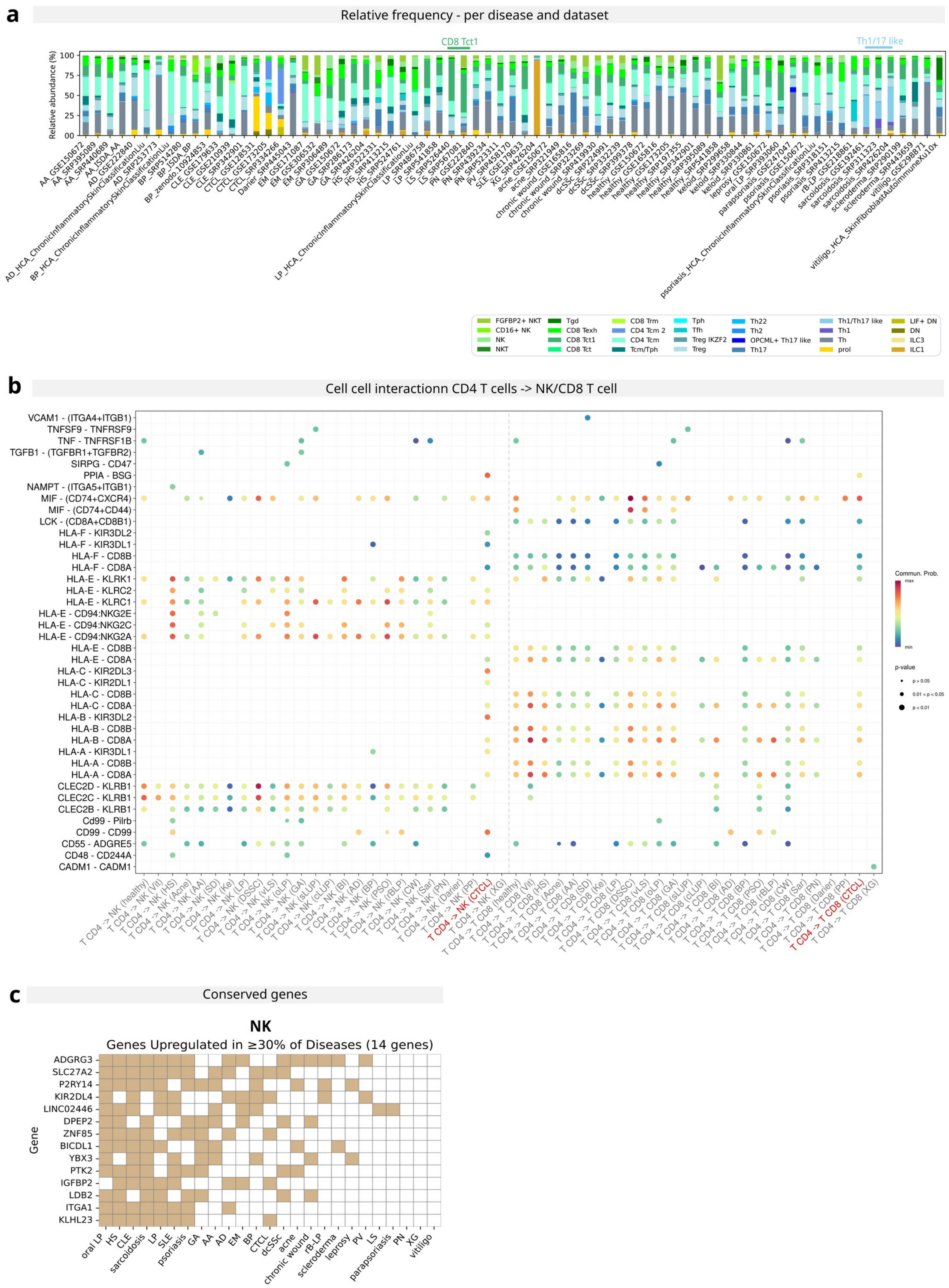


***a)** Cell Type Frequency by Disease and Dataset: Relative abundance of cell types across diseases and datasets, emphasizing the conserved high frequency of a specific T cell subtype in certain diseases. **b)** Cell–Cell Interaction Network: Bubble plot depicting interactions between CD4 T cells and NK cells (left) and between CD4 T cells and CD8 T cells (right). **c)** Conserved Differential Gene Expression: Top percentage of conserved upregulated genes in disease compared to healthy controls ($log_2$ fold change > 2, adjusted p-value < 0.5).*

***Supplementary Figure 5: Fibroblasts***

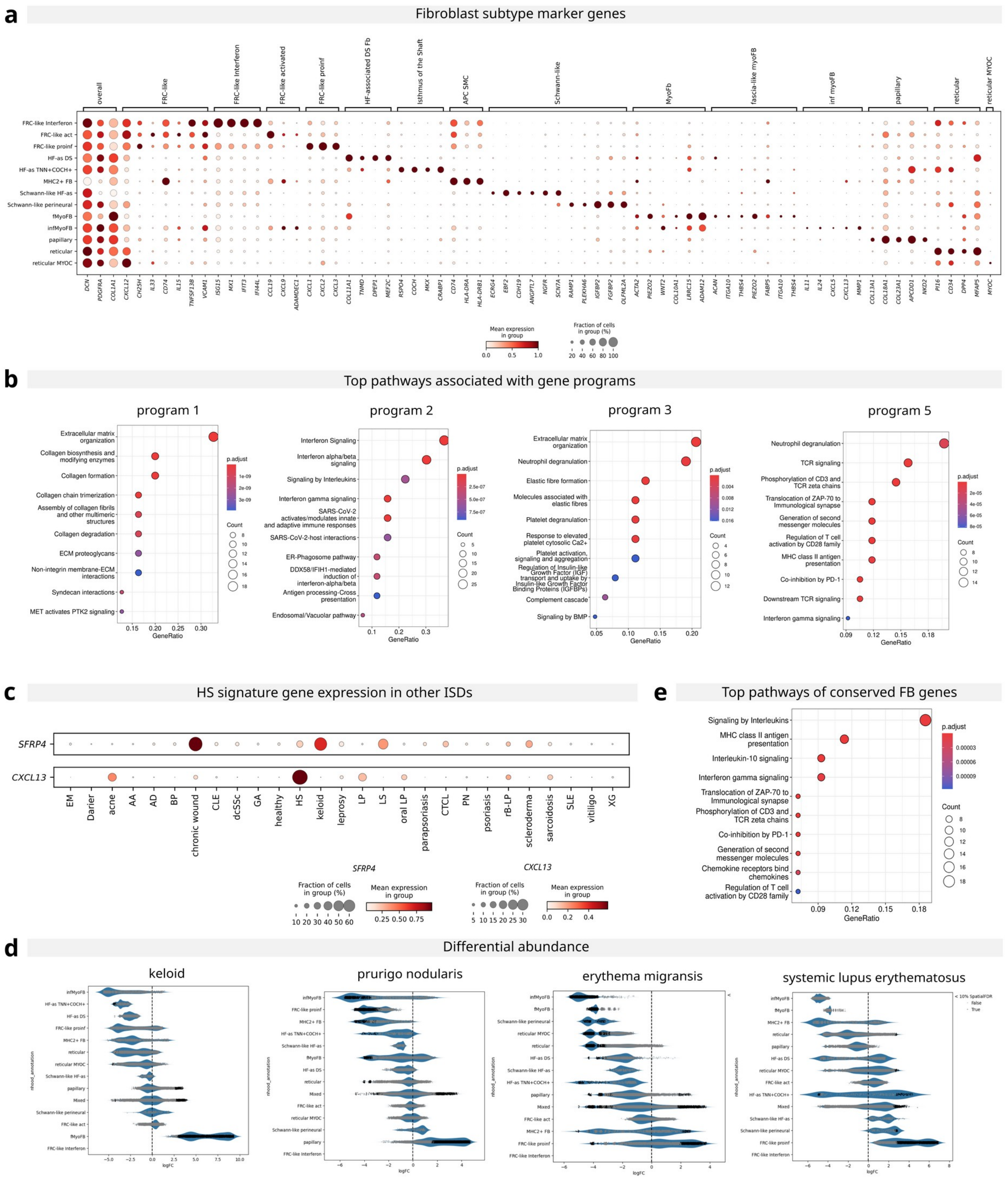


***a)** Marker Gene Expression in Fibroblast Subsets: Dot plot showing the expression of marker genes across fibroblast subsets. **b)** Pathway Enrichment Analysis: Top-ranked pathways associated with selected gene programs in fibroblasts. **c)** Disease-Specific Gene Expression: Expression levels of CXCL13 and SFRP4 in fibroblasts across diseases. **d)** Differential Abundance of Fibroblast Neighborhoods: Beeswarm plot of differential abundance testing for selected diseases. Each dot represents a neighborhood of cells, with significantly enriched neighborhoods (FDR < 10%) highlighted in black. **e)** Conserved Pathway Analysis: Top pathways derived from conserved fibroblast genes (present in >50% of ISDs).*

***Supplementary Figure 6: Schwann cells and mural cells***

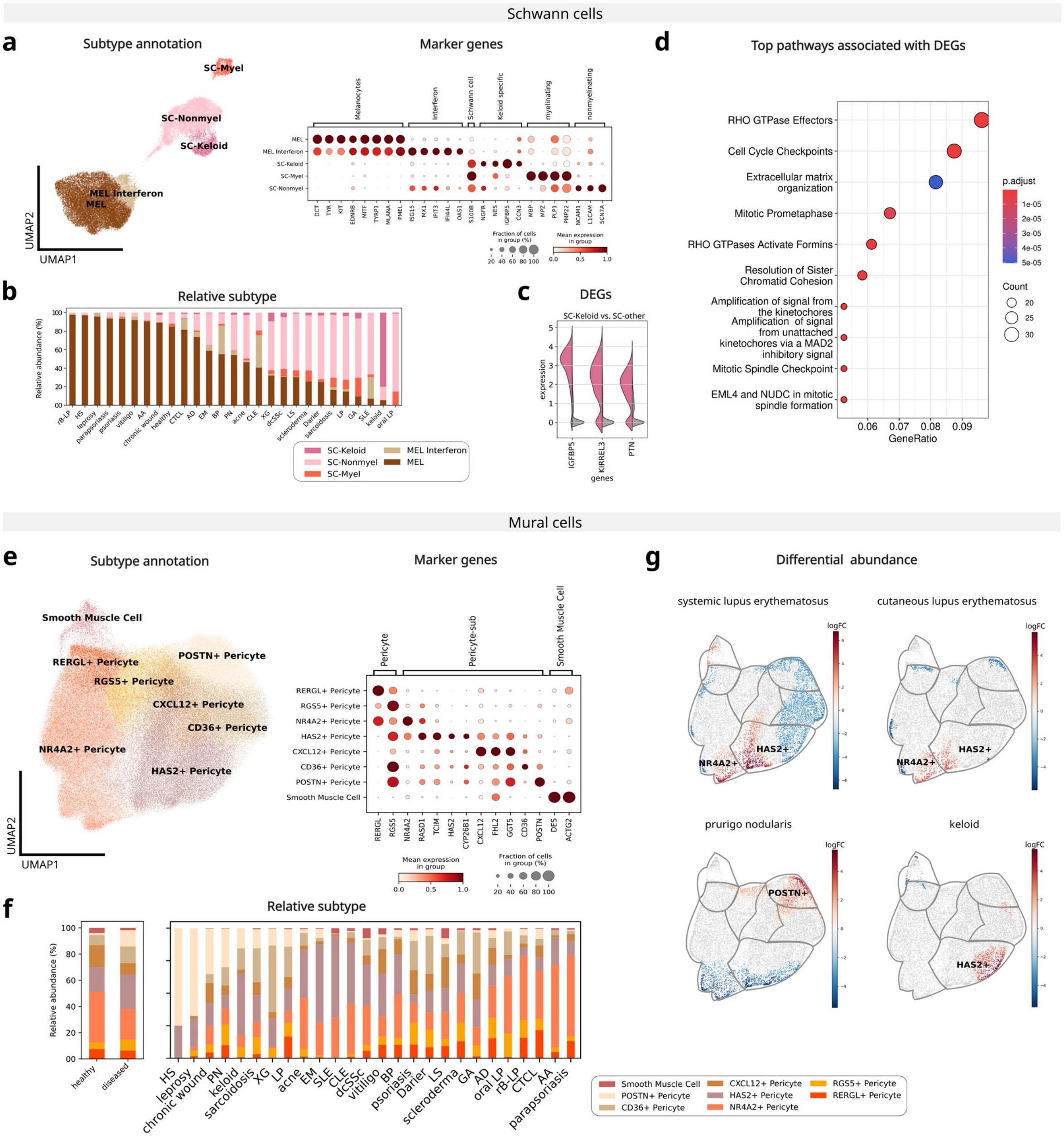


***a)*** *Subtype Annotation and Marker Expression: UMAP embedding of melanocyte and Schwann cell subtypes, accompanied by a dot plot of corresponding marker gene expression.* ***b)*** *Cell Subset Distribution by Disease: Relative abundance of melanocyte and schwann cell subsets across diseases.* ***c)*** *Differential Gene Expression in Schwann Cells: DEGs distinguishing keloid-associated schwann cells from other subsets.* ***d)*** *Pathway Enrichment in Schwann Cells: Top-ranked pathways enriched in unique DEGs of schwann cells compared to healthy skin.* ***e)*** *Subtype Annotation and Marker Expression: UMAP embedding of mural cell subtypes with a corresponding marker gene dot plot.* ***f)*** *Mural Cell Subset Distribution: Relative abundance of mural cell subsets in healthy samples and diseased samples, as well as stratified by disease.* ***g)*** *Differential Abundance of Mural Cell Neighborhoods: Differential abundance testing for selected diseases.*

***Supplementary Figure 7: Keratinocytes***

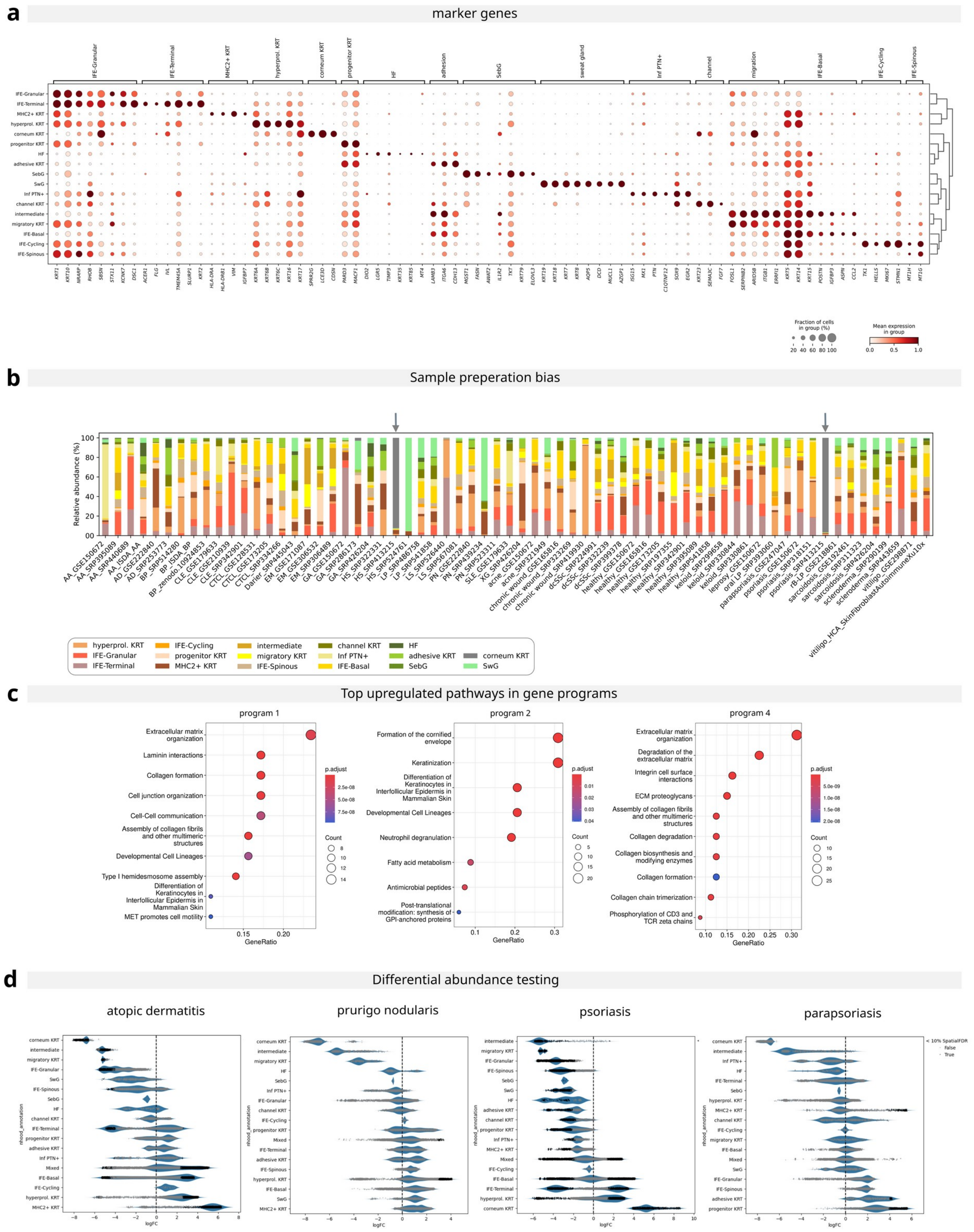


*a) Marker Gene Expression in Keratinocytes: Dot plot illustrating the expression of marker genes across keratinocyte subtypes. b) Cell Type Frequency by Disease and Dataset: Relative abundance of keratinocyte subtypes, stratified by disease and dataset, with emphasis on dataset-specific biases in cell composition. c) Pathway Enrichment Analysis: Top pathways derived from gene modules in keratinocytes. d) Differential Abundance of Keratinocyte Neighborhoods: Beeswarm plot of differential abundance testing for selected diseases. Each dot represents a neighborhood of cells, with significantly enriched neighborhoods (FDR < 10%) highlighted in black.*

**_Supplementary Figure 8: Myeloid cells_**

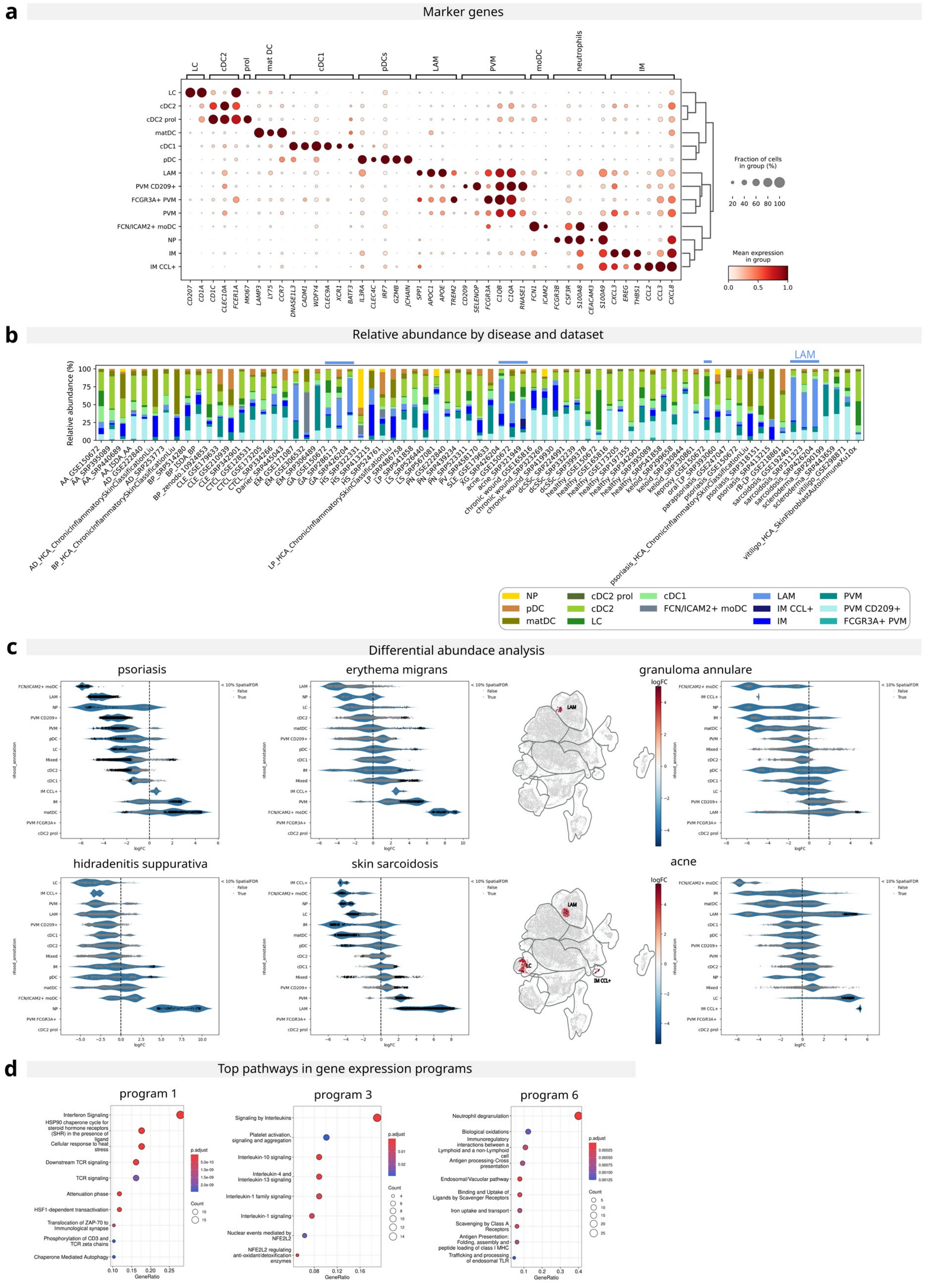


*__a)__ Marker Gene Expression in Myeloid Cells: Dot plot showing the expression of marker genes across myeloid cell subtypes.* *__b)__ Myeloid Cell Frequency by Disease and Dataset: Relative abundance of myeloid subtypes, stratified by disease and dataset, highlighting conserved high frequencies of lipid associated macrophages in granulomatous diseases and acne.* ***c)***

*Differential Abundance of Myeloid Neighborhoods: Beeswarm and UMAP plots of differential abundance testing for selected diseases. Each dot in the beeswarm plot represents a neighborhood of cells, with significantly enriched neighborhoods (FDR < 10%) highlighted in black.* ***d)*** *Pathway Enrichment in Myeloid Cells: Top pathways derived from myeloid gene modules.*

***Supplementary Figure 9: Endothelial cells***

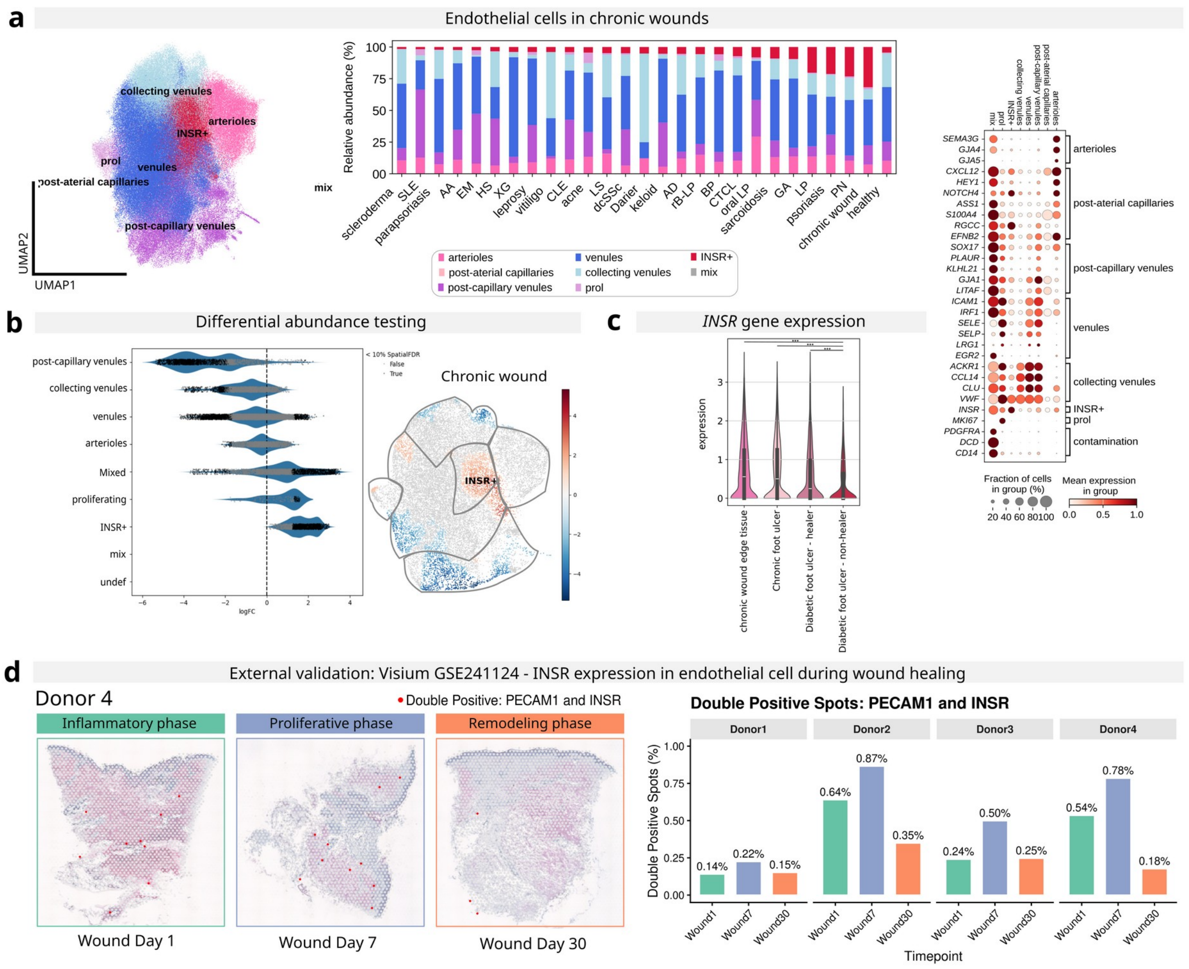


***a)*** *Endothelial Cell Subtype Characterization: UMAP embedding of endothelial cell subtypes, relative cell type distribution across diseases, and a dot plot of marker gene expression.* ***b)*** *Differential Abundance in Chronic Wounds: UMAP and beeswarm plot of differential abundance testing for chronic wounds. Each dot represents a neighborhood of cells, with significantly enriched neighborhoods (FDR < 10%) highlighted in black.* ***c)*** *INSR Expression in Chronic Wound Subsets: Expression levels of INSR across endothelial cell subsets in chronic wounds.* ***d)*** *External Validation in Spatial Data (GSE241124): Representative hematoxylin and eosin (H&E) stained sections depicting the distinct wound healing phases in donor 4. PECAM1/INSR double-positive spots are highlighted in green. The accompanying bar plot quantifies the relative frequency of double-positive spots across donors and time points.*

### Extended Data

***Extended Data 1 - Harmonized cell annotation through the Healthy Skin Atlas***

To achieve robust cell-type annotation of specialized subpopulations, we leveraged marker gene profiles from the Healthy Skin Cell Atlas (HSCA) and validated annotations via integration between HSCA and our Inflammatory Skin Disease Atlas (ISDA). Mapping examples for fibroblasts and mural cells are shown in Extended Data Fig. 1a,b. Highlighting in both cell types that disease specific subtypes can be found.

***Extended Data Fig. 1: Mapping of inflammatory skin disease atlas to  healthy skin cell atlas***

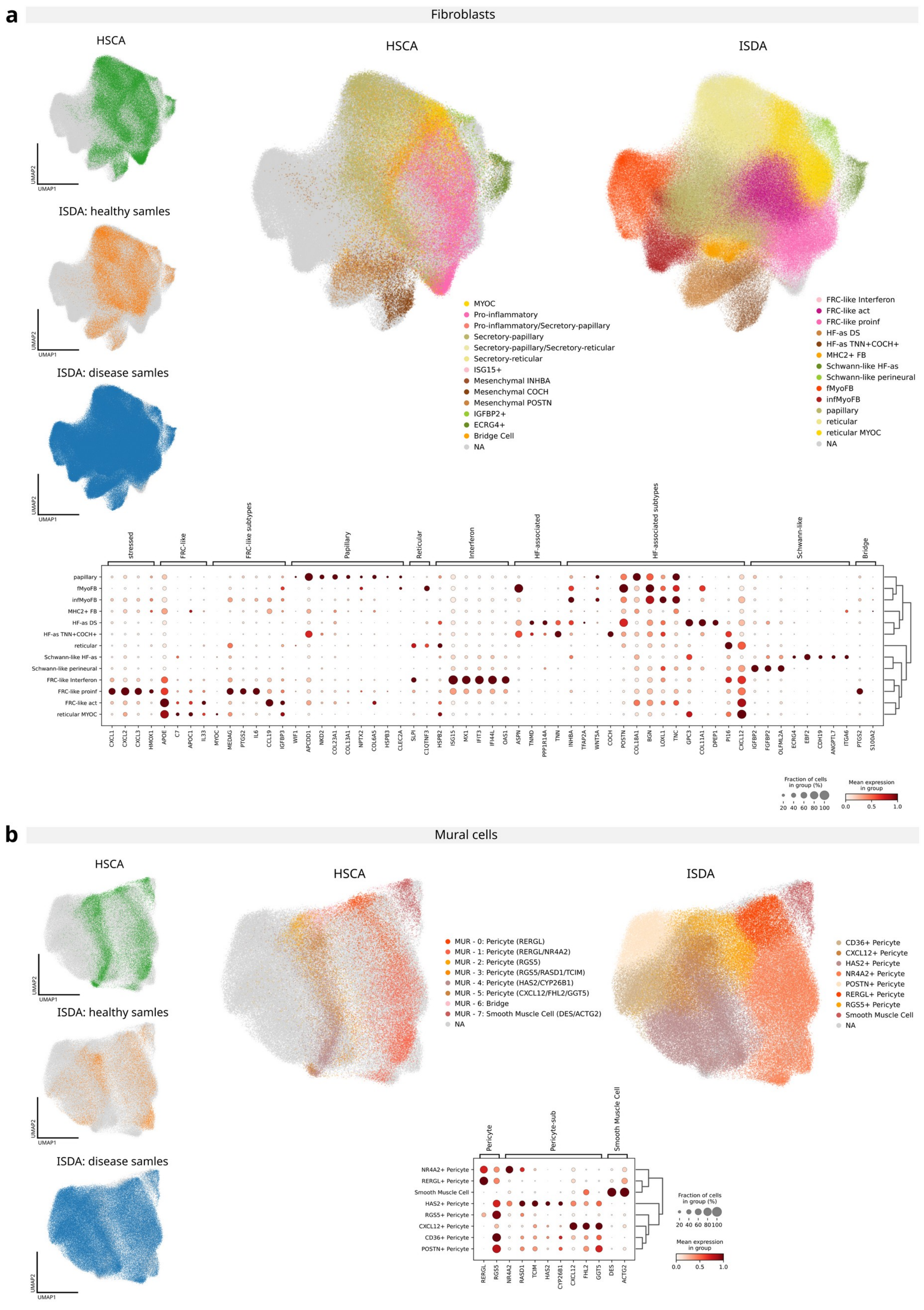


*a,b) Integrated UMAP Embeddings of HSCA and ISDA Datasets:. Highlighting: HSCA cells (left top); healthy ISDA cells (left middle); disease ISDA cells.; cell annotation of HSCA cells (middle), cell annotation ISDA cells (right); HSCA marker gene expression on ISDA cell annotation (bottom). HSCA, healthy skin cell atlas; ISDA, inflammatory skin disease atlas*

***Extended Data 2 - Technical bias through sampling and bioinformatics methods***

Sample preparation and data processing are critical steps in single-cell analyses, requiring careful consideration to minimize bias and ensure robust interpretation of results. Integrating this large amount of data enabled us to directly compare their influence on the results. During quality check of individual dataset we observed that a sample from GSE165816 had gene ID artifacts (dates embedded in identifiers; Extended Data Fig. 2a), a known issue in public datasets[1], underscores the critical need for rigorous preprocessing in meta-analyses. Apart from a higher mitochondrial content in Seqwell S3[2] samples, we did not observe major differences in choice of technology based on standard quality matrixes (Extended Fig. 2b). Two of the included datasets (GSE150672 and GSE17087) were aligned to the Human Reference Genome version 19 (Hg19), whereas all other datasets were aligned to Hg38 according to the respective methods. We deliberately included these datasets in order to assess the effect of different reference genomes. As expected, we observed that Hg19 aligned datasets used previous HGNC-approved nomenclature[3] (Extended Fig. 2c). Surprisingly, this was also true for two further datasets (HCA_SkinFibroblastAutoimmuneXu10x and GSE128531) which were supposedly aligned to Hg38. This highlights the importance of consistent processing to prevent misinterpretations. Any findings in the downstream analysis representing the genome bias were excluded.

Sampling strategies introduced marked variability in cell type composition across diseases. For instance, lichen sclerosus samples showed diversity probably linked to sex and biopsy site (labia minora/interlabial sulcus vs. foreskin), while dataset variations in atopic dermatitis, lichen planus and scleroderma are likely linked to the usage of fresh vs frozen samples, which has a known impact on cell type recovery and especially keratinocytes[4]. Sarcoidosis samples from GSE192461 showed higher macrophage frequencies. Here, the authors first sorted for CD45+ and CD45- negative cells and then pooled them in an equal ratio[5]. Vitiligo samples from HCA_SkinFibAutoimmuneXu10x contained melanocytes, contrasting with disease hallmarks. Here, the authors took samples at the edge of lesions[6] (Extended Fig. 3a). We did not observe major bias on cell type composition concerning bodysite location

(Extended Fig. 3b). Moreover, we observed a dataset-specific keratinocyte cluster that uniquely expressed corneum-associated genes like SPRR2G and LCE3D (Extended Fig. 3c, Supplementary Fig. 1c). These keratinocytes were found in two diseases (psoriasis and hidradenitis suppurativa), yet originated from the same dataset. Here, the authors analysed cells emigrating from the tissue after 48 hours of incubation[7], which may have resulted in a processing-related artefact. This dataset was excluded from subsequent keratinocyte-related analyses. CLE and SLE samples showed high cell type variance (Extended Fig. 3d). Here, the authors split samples into dermis and epidermis[8], reflected in different cell type compositions[9]. We also observed biological variation for diseases within datasets: xanthogranuloma, parapsoriasis, and leprosy displayed heterogeneity between samples and also for chronic wound, acne, bullous pemphigoid, alopecia areata, erythema migrans and CTCL heterogeneity seemed to be dataset independent (Extended Fig. 3a,d). Whereas oral lichen planus, rB-LP, pemphigus vulgaris and Darier disease samples appeared homogeneous (albeit single-dataset limited) and keloid and dcSSc appeared homogeneous even across datasets. Overall no major differences were observed based on sequencing technology but rather sampling strategy. These findings underscore the critical influence of sample preparation on cell type composition, a factor that must be considered when interpreting downstream results. Nevertheless, in all diseases cell type composition reflects known pathology, such as higher B cell abundance in HS.

*Extended Data Fig. 2: Dataset bias based on data processing*

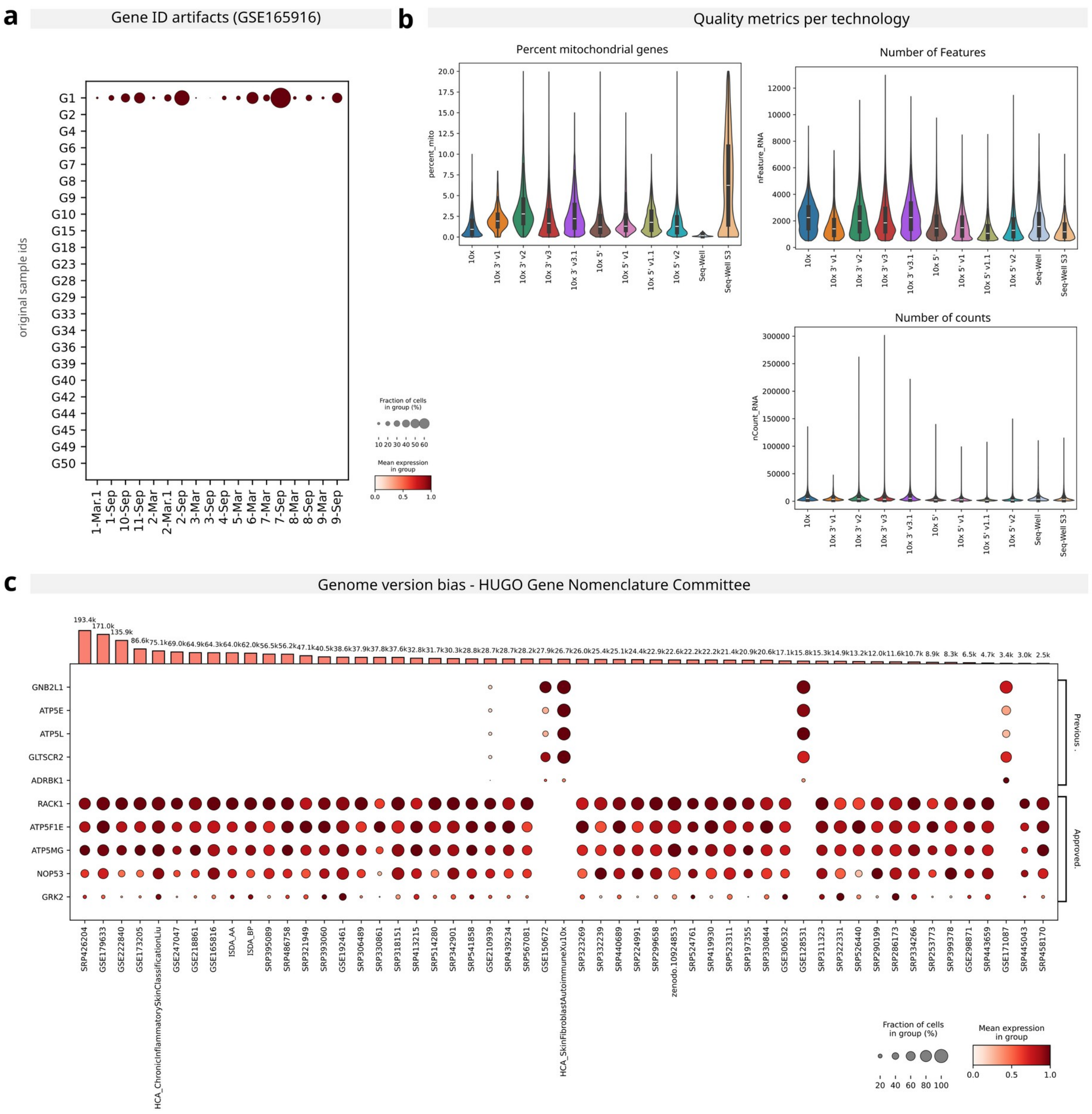


***a)*** *Gene ID Artifacts: Dot plot highlighting genes artefacts in one dataset.* ***b)*** *Quality Metrics by Sequencing Platform: Violin plots comparing key quality metrics across sequencing technology.* ***c)*** *Genome Version Bias: Dot plot illustrating bias related to genome version. Top: Bar plot showing cell counts per dataset.*

***Extended Data Fig. 3: Dataset bias based on sampling strategy***

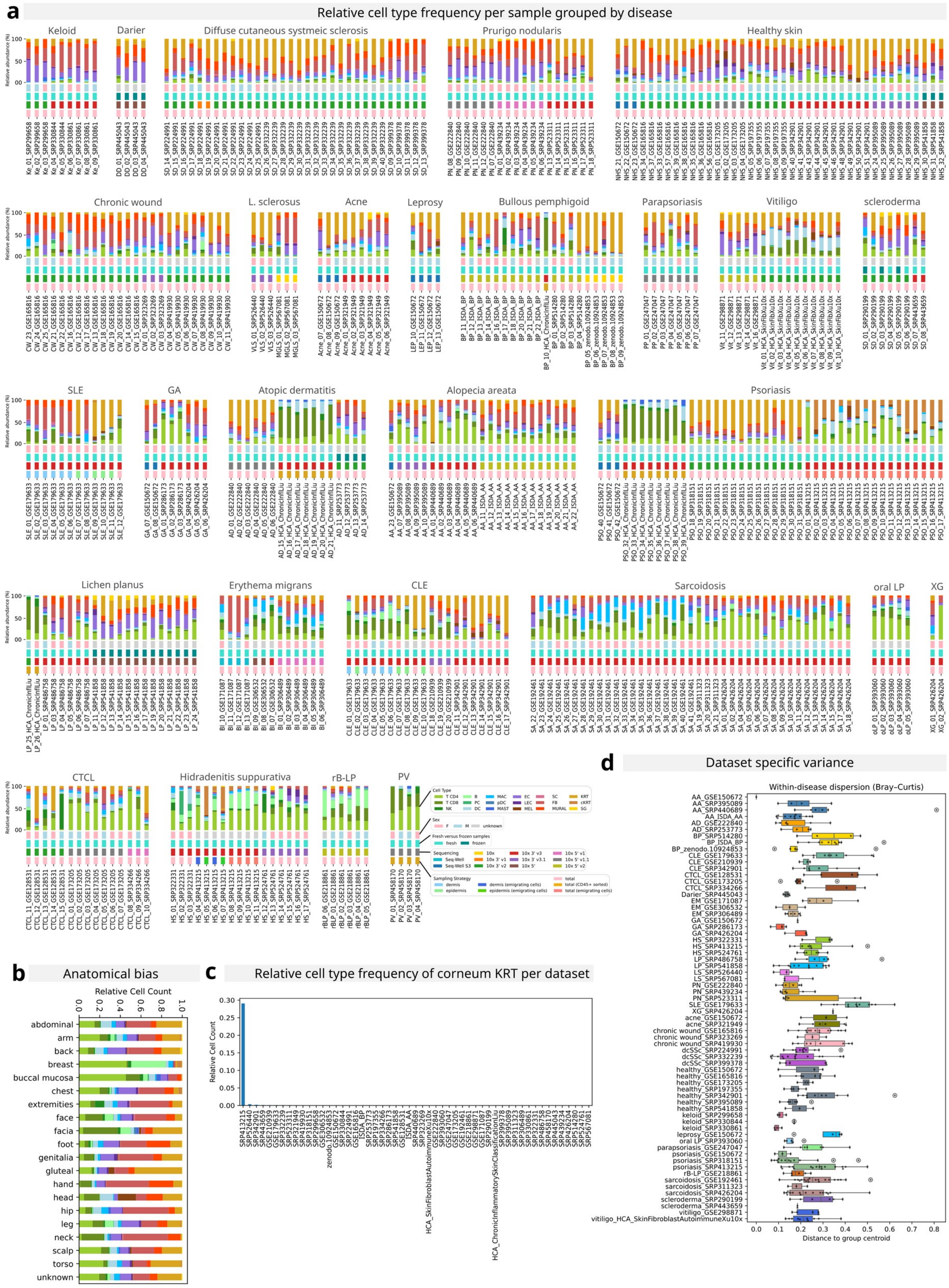


*a) Sample Metadata Overview: Stacked bar plots showing the relative frequency per sample, grouped by disease and stratified as follows (top to bottom): Cell type composition, Sex distribution, Fresh vs. frozen sample status, Sequencing technology, Sampling strategy. **b)** Body site bias: Stacked bar plots showing the relative frequency per anatolmical region stratified by cell type. **c)** Dataset-Specific Keratin Bias: Highlighting the corneum KRT gene bias originating from a single dataset. **d)** Cellular Composition Diversity: Diversity index quantifying differences in cellular composition* *between samples within each disease and dataset. Note: Samples containing CD45+-sorted cells were excluded from analyses in panel b,c).*

#### *Extended Data only references*
